# Characterization of covalent inhibitors that disrupt the interaction between the tandem SH2 domains of SYK and FCER1G phospho-ITAM

**DOI:** 10.1101/2023.07.28.551026

**Authors:** Frances M. Bashore, Vittorio L. Katis, Yuhong Du, Arunima Sikdar, Dongxue Wang, William J. Bradshaw, Karolina A. Rygiel, Tina M. Leisner, Rod Chalk, Swati Mishra, Andrew C. Williams, Opher Gileadi, Paul E. Brennan, Jesse C. Wiley, Jake Gockley, Gregory A. Cary, Gregory W. Carter, Jessica E. Young, Kenneth H. Pearce, Haian Fu, the Emory-Sage-SGC TREAT-AD Center, Alison D. Axtman

## Abstract

RNA sequencing and genetic data support spleen tyrosine kinase (SYK) and high affinity immunoglobulin epsilon receptor subunit gamma (FCER1G) as putative targets to be modulated for Alzheimer’s disease (AD) therapy. FCER1G is a component of Fc receptor complexes that contain an immunoreceptor tyrosine-based activation motif (ITAM). SYK interacts with the Fc receptor by binding to doubly phosphorylated ITAM (p-ITAM) via its two tandem SH2 domains (SYK-tSH2). Interaction of the FCER1G p-ITAM with SYK-tSH2 enables SYK activation via phosphorylation. Since SYK activation is reported to exacerbate AD pathology, we hypothesized that disruption of this interaction would be beneficial for AD patients. Herein, we developed biochemical and biophysical assays to enable the discovery of small molecules that perturb the interaction between the FCER1G p-ITAM and SYK-tSH2. We identified two distinct chemotypes using a high-throughput screen (HTS) and orthogonally assessed their binding. Both chemotypes covalently modify SYK-tSH2 and inhibit its interaction with FCER1G p-ITAM.

## Introduction

SYK is a key signaling kinase in adaptive and innate immunity. The 72 kDa protein is composed of a catalytic kinase domain, a flexible interdomain region B (IB), and two tandem SH2 (SYK-tSH2) domains connected by interdomain A (IA) (Figure 1A) (1). The tandem SH2 domains of SYK bind to phosphorylated ITAM motifs such as the one on FCER1G within Fc receptor complexes (FceRI, FcgRI, and FcgRIIIa) (2, 3).

**Figure 1.**
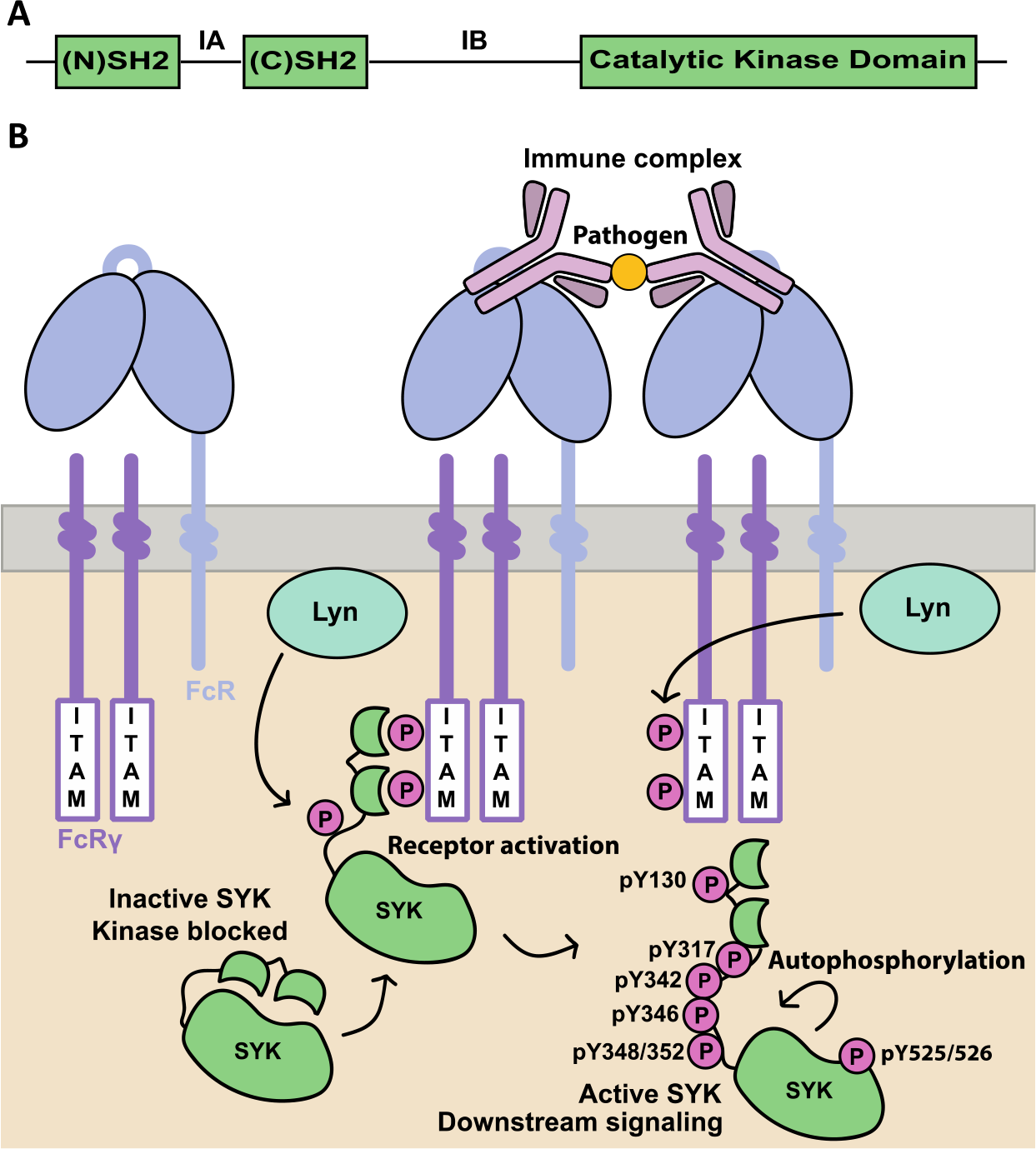
SYK topology and activation mechanism with Fc receptors. **(A)** Domain representation of SYK. (N)SH2 domain (residues 8–118), interdomain A (IA), (C)SH2 domain (residues 163–264), interdomain B (IB), and the catalytic kinase domain (364–620). **(B)** The activation mechanisms of autoinhibited SYK via both autophosphorylation and engaging p-ITAM on Fc receptors. Tyrosines phosphorylated via these mechanisms are highlighted.

Generation of the doubly phosphorylated ITAM (p-ITAM) is initiated by the stimulation of receptor complexes with IgE, IgG or IgM, which triggers the phosphorylation of ITAM motifs by Lyn or Fyn Src kinases (Figure 1B) (4). SYK binds to the p-ITAM sequence region on the γ chain within Fc receptor complexes via the SYK-tSH2 domains (2). Subsequent Lyn-mediated phosphorylation of the SYK SH2 interdomain linker region at residues Tyr348/352 leads to SYK activation (5). Phosphorylation of the SYK activation loop at residues Tyr525/526 is reported to result in further SYK activation, as well as additional SYK autophosphorylation or transphosphorylation and direct phosphorylation by Src family kinases at Tyr residues in its interdomain A and B, kinase, and C-terminus regions (1, 6, 7, 8). Engagement of SYK-tSH2 domains with p-ITAM releases the kinase domain from its autoinhibited state by releasing its interaction with the catalytic kinase domain (5). SYK is not limited to Fc receptor complex interaction, as it also plays a significant role in both B cell receptor (BCR) and triggering receptor expressed on myeloid cells 2 (TREM2) signaling (9, 10, 11). Broadly, SYK activation can initiate downstream signaling pathways such as PI3K, AKT, ERK, mTOR, NLRP3 inflammasome, among others (12, 13, 14). These signaling pathways are proposed to play essential roles in driving AD pathology and therefore chemical probes to inhibit SYK activation are of therapeutic relevance (15, 16, 17).

Numerous ATP-competitive inhibitors that bind to the SYK kinase domain have been developed to date. These inhibitors do not disrupt the interaction between the SYK-tSH2 domain and p-ITAM on Fc receptors. ATP-competitive SYK inhibitors, entospletinib, BAY61-3606, PRT062607, and TAK-659, are potent and moderately selective (18, 19, 20, 21). Additional potent SYK inhibitors such as R406 are less selective, as they also potently inhibit multiple other kinases (22, 23). Fostamatinib is the prodrug of R406 that was approved by the FDA for the treatment of chronic immune thrombocytopenia (24). The small molecule inhibitors entospletinib, TAK-659, and cerdulatinib are in clinical trials for various cancers (25). SYK inhibitors that exhibit good selectivity for SYK over other kinases represent useful tool compounds to interrogate SYK-mediated biology. For example, BAY61-3606 treatment or SYK knockdown with shRNA-SYK reduces phosphorylated and total tau levels *in vitro* (SH-SY5Y cells, 5 μM) and reduces p-AKT, p-mTOR, and p-p70 S6K levels (26). Furthermore, *in vivo* mouse models of AD (Tg Tau P301S mice) showed reduced levels of phospho-tau (p-tau) and total tau (t-tau) after SYK inhibition with BAY61-3606 (20 mg/kg) over a course of 12 weeks (26). Despite this, ATP-competitive SYK inhibitors are likely to have only limited selectivity across the human kinome and may not be suited for development as drugs for treating a chronic neurodegenerative disease such as AD. Less research has focused on inhibitors that bind to the SYK-tSH2 domain and/or that inhibit p-ITAM binding to SYK. A covalent inhibitor (**1**) and cysteine oxidation with hydrogen peroxide was reported to disrupt the binding of SYK-tSH2 domains with a doubly phosphorylated ITAM from a T-cell antigen receptor ζ-chain in a fluorescence polarization inhibition assay. Compound **1** binds to all 4 of the cysteines available on SYK-tSH2 as determined by mass spectrometry (27). Cys206 was identified by the authors as the key residue for inhibition of p-ITAM binding to SYK-tSH2 domains with **1**. Mutation of Cys206 to either a serine or alanine residue resulted in a complete loss of inhibition by **1**. Cys206 is located within the (C)SH2 domain and covalent binding of this residue by **1** presumably inhibits p-ITAM binding allosterically. The authors suggest oxidation of cysteine residues in the SYK-tSH2 domain region could be a viable strategy to regulate cell signaling through SYK inhibition.

Inhibition or genetic manipulation of SYK demonstrates its importance in AD. SYK can be activated in response to Aβ exposure and in turn SYK activation also correlates with increased Aβ and tau hyperphosphorylation (28). Treatment with a SYK inhibitor (BAY61-3606) in mouse tauopathy models resulted in decreased tau accumulation, neuronal and synaptic loss, neuroinflammation, and reversed defective autophagy (15). Additionally, SYK knockdown via shRNA mimicked pharmacological inhibition with SYK inhibitor BAY61-3606 in SH-SY5Y cells, resulting in a decrease in p-tau and t-tau levels. Inhibition of SYK with BAY61-3606 prevented lipopolysaccharide (LPS)-induced neuronal loss in primary neuron-glia cultures, and reduced microglial-mediated phagocytosis of synapses and neurons (29). The observed decrease in microglial phagocytosis, resulting from SYK inhibition, led the authors to propose that it was the probable cause of neuroprotection. Conversely, SYK is also reported to be neuroprotective in microglia with SYK deletion causing increased Aβ in mouse models of AD (16). Phagocytosis of Aβ by microglia was reduced in 5×FAD mice with SYK deletion and it is proposed that the dysregulation of GSK3β in the absence of SYK could be contributing to the decreased phagocytosis observed. These data suggest that further interrogating the role of SYK in AD pathology is of therapeutic relevance. FCER1G is reported to play roles in phagocytosis, microglial activation, and inflammation, which are also processes that are aberrant in AD (30, 31). FCER1G has been previously identified as a risk gene for AD in genome-wide association studies (GWAS), and SYK was identified as being differentially expressed in an APP/PS1 (APPtg) mouse model in an age and genotype comparison despite being statistically insignificant in published GWAS (32).

Herein, we used several biochemical and biophysical assays that enable screening for small molecules that inhibit the interaction of SYK-tSH2 and the FCER1G p-ITAM. As experimental validation for pursuing the target, we first characterized the expression levels of SYK in the brain of mice in various AD models using a validated SYK antibody and examine the expression levels of SYK in human induced pluripotent stem cell (hiPSC) derived neurons and microglia. We also developed a time-resolved fluorescence energy transfer (TR-FRET) assay and a miniaturized TR-FRET assay for an ultra-high-throughput screen (uHTS) campaign to identify protein–protein interaction (PPI) disruptors. The uHTS of the Emory chemical diversity compound library, containing 138,214 compounds, identified three putative hit compounds from two orthogonal chemical series. A secondary biophysical assay, bio-layer interferometry (BLI), was used to confirm direct binding to SYK-tSH2. Intact mass analyses helped identify the active compound in one of two chemical series and confirm that all three compounds covalently modify SYK-tSH2. Importantly, these small molecules inhibit the interaction of full length SYK and FCER1G through an orthogonal chromatography-based GST pull-down (GST-PD) assay in cell lysates. We hypothesize the covalent modification of cysteine residues could be a viable strategy to disrupt the interaction of FCER1G and SYK. This strategy could help to interrogate the importance of this interaction for AD pathology.

## Results

### Bioinformatics identifies SYK and FCER1G as putative targets for AD and mRNA expression is confirmed in microglia

To prioritize putative target proteins for AD, the Emory-Sage-SGC TREAT-AD Center has developed a Target Risk Score (TRS) to prioritize putative target proteins for AD (Figure 2A–B) (33) and represents a composite of genetic and omic dimensions of risk. The genetic risk metric uses evidence from genetic association studies (GWAS, GWAX, and QTL), variant severity analysis, and phenotypic evidence from humans and model organisms. The multi-omic risk metric is based on signatures of differential expression from meta-analyses of transcriptomic and proteomic data from the AMP-AD consortium.

**Figure 2.**
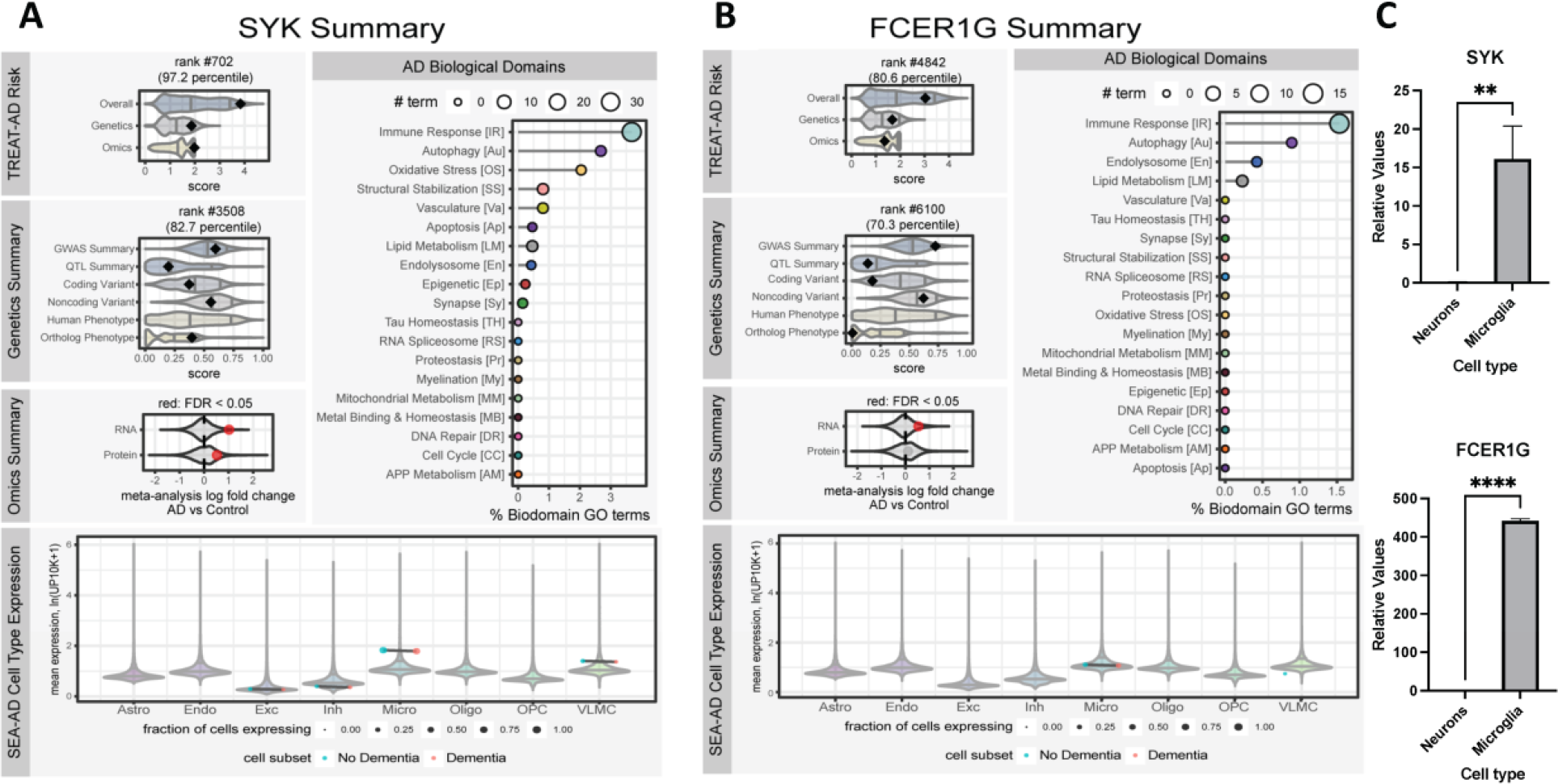
TREAT-AD bioinformatic summaries for SYK **(A)** and FCER1G **(B)**. For each target, the composite Target Risk Score (Overall, top left) is a sum of the Genetics and Omics risk dimensions. The Genetics Summary includes the average rank of gene-level significance values from GWAS and QTL studies, variant severity analysis, and phenotype summaries for both human genes (Hsap) and orthologous genes. The Omics Summary represents the results from meta-analyses of AMP-AD transcriptomic and proteomic datasets showing the effect size (log fold change) and significance (red points are detected at an FDR < 0.05). For all plots no point plotted indicates that the gene was not scored or measured in that dimension. The AD Biological Domains shows the fraction of all terms annotated to a biological domain to which the gene is annotated and the size of the point indicates the number of terms. The SEA-AD Cell Type Expression panel shows the mean expression values for the gene per cell subclass, stratified by cognitive status of the donor, where expression is measured as the natural log[[number of unique molecular identifiers (UMIs) for each gene in a given cell divided by the total number of UMIs in the same cell divided by 10,000] plus 1]. The size of each point represents the fraction of cells from each cell type expressing the gene. **(C)** mRNA expression levels of SYK and FCER1G in hiPSC-derived neurons and microglia.

The TRS for SYK is 3.84, which places SYK in the 97^th^ percentile of all scored targets. SYK is found to be differentially expressed in brains of AD patients relative to controls in both transcriptomic and proteomic studies, giving it an omics score of 1.98 out of 2. The genetic contribution to the SYK TRS is 1.86 out of 3 (82^nd^ percentile), which is based on modest and consistent scores from the association studies (GWAS and QTL) as well as variant severity analyses, as well as a high correspondence of phenotypes attributable to SYK orthologs in model organisms and AD related phenotypes.

The TRS for FCER1G is slightly lower than that for SYK at 3.03 (80^th^ percentile). FCER1G is only found to be significantly differentially expressed based on meta-analyses of transcriptomic datasets, leading to a slightly lower omics risk score of 1.36 out of 2. The genetics risk computed for FCER1G is also less than SYK — 1.67 out of 3 — despite having a relatively higher GWAS summary metric, owing in large part to a lower contribution from coding variant severity and a lower correspondence between FCER1G ortholog phenotypes and AD.

The Emory-Sage-SGC TREAT-AD Center has also developed the Biological Domains of AD as an approach to objectively define and categorize disease relevant risk. There are 19 distinct biological domains derived from various molecular endophenotypes that have been linked with AD, which are defined functionally by largely non-overlapping sets of Gene Ontology (GO) terms. Both SYK and FCER1G have the strongest annotation to the Immune Response and Autophagy biological domains. Finally, to understand the relevant cell types for each gene, summary expression from the SEA-AD snRNA-seq dataset was considered (34). Expression of both SYK and FCER1G is detected broadly in microglia. The mRNA levels of SYK and FCER1G were measured in hiPSC derived neurons and microglia (Figure 2C). Significantly higher levels of both SYK and FCER1G were quantified in microglia compared to neurons, which is consistent with the reported roles of the activation of SYK and FcγR in microglia (35).

### Identification of covalent inhibitors of the interaction between FCER1G p-ITAM and SYK-tSH2 via HTS

To facilitate identification of inhibitors of the SYK-FCER1G interaction, we developed a TR-FRET assay using recombinant protein comprising the tSH2 domain of SYK (residues M6-N269) fused to an N-terminal 6×His tag (Figure S1A). The TR-FRET assay is based on the interaction of a FITC-labeled FCER1G p-ITAM peptide to the 6×His-SYK-tSH2 protein in complex with a terbium-conjugated anti-His antibody (Tb-Ab) (Figure 3B). Once excited, a TR-FRET signal is generated from the Tb donor to the FITC acceptor fluorophore. The TR-FRET signal produced by this interaction can be competed in a dose-dependent manner by the addition of unlabeled FCER1G p-ITAM peptide (IC_50_ = 12 nM). As expected, the non-phosphorylated FCER1G ITAM peptide has no effect on the TR-FRET signal (IC_50_ >1000 nM, Figure 3C). To enable uHTS for large scale screening of small molecule libraries, we miniaturized the TR-FRET assay from 96-well plate format to a 1536-well. A primary screen of 138,214 small molecules from the Emory Chemical Biology Discovery Center (ECBDC) identified 294 primary positives in which the signal was less than 60% of the control, corresponding to a hit rate of 0.2% (Figure 3D). The 294 putative hits were confirmed in a dose-response format from the library stock plates and 135 compounds were confirmed to have an IC_50_ <20 μM. Of these 135 small molecules, 112 were commercially available for re-evaluation in the primary TR-FRET assay, leading to confirmation of 54 compounds with IC_50_ <30 μM. This set of 54 potential leads was counter-screened in a TR-FRET assay developed for an unrelated PPI (moesin-CD44) with the same donor and acceptor fluorophores in order to exclude fluorophore interference compounds (36, 37). After removing pan-assay interference compounds (PAINS) and cross-reactive compounds from the moesin-CD44 assay, 40 compounds were progressed for further hit validation in a panel of secondary assays. These compounds were triaged first in a GST-PD assays with full length SYK and FCER1G, followed by confirmation in a biophysical assay to determine direct binding.

**Figure 3.**
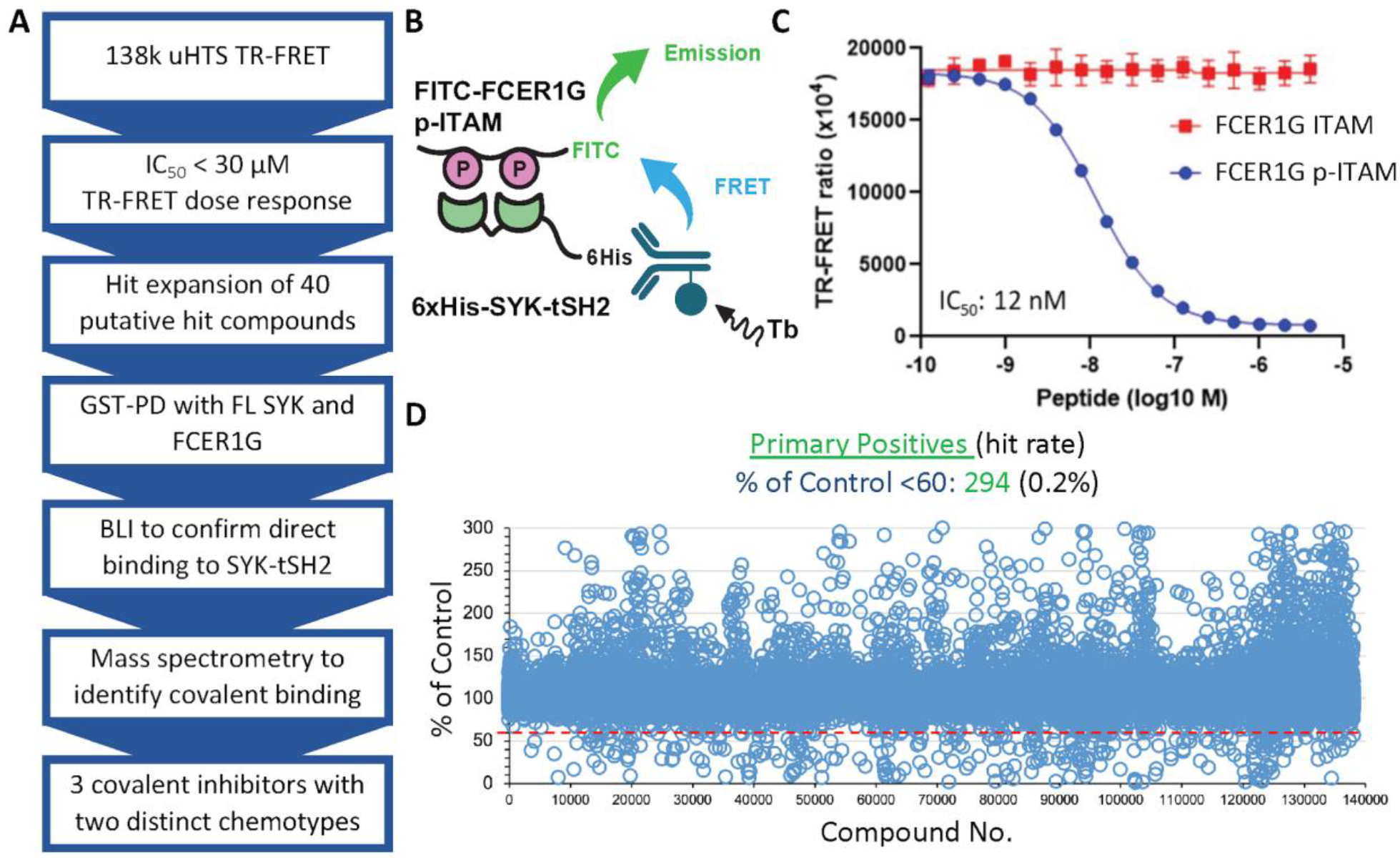
Compound screening cascade and the development and use of a biochemical TR-FRET assay to evaluate inhibition of the PPI between SYK-tSH2 and the FCERG1G p-ITAM peptide. **(A)** Compound screening cascade. **(B)** and **(C)** TR-FRET assay with 6xHis-SYK-tSH2 and a p-ITAM-FITC tagged peptide. Unlabeled FCER1G p-ITAM can displace p-ITAM-FITC, while FCER1G ITAM does not. **(D)** Primary uHTS of 138,214 compounds with 294 primary positive identified.

Pull-down assays were used as an orthogonal assay to determine if the compounds could disrupt the PPI between full length SYK and FCER1G proteins from cell lysates. To optimize GST pulldown assays, full length GST-SYK and FCER1G-Flag were co-expressed in HEK293 cells prior to lysis and the protein expression and cell lysate concentrations tested (Figure 4A). All 40 compounds that progressed from uHTS were tested via GST-PD at a single concentration (100 μM) to determine if they could inhibit the PPI between the full-length proteins, with the p-ITAM FCER1G peptide used as a positive control (Figure 4B). Among the 40 compounds tested for PPI disruption at 100 μM, 11 compounds inhibited the primary GST-PD at 100 μM, and 5 of these 11 compounds (**4, 13, 24, 32**, and **37**) were confirmed to disrupt the PPI in a dose-response format. Compounds **13, 24**, and **37** showed the greatest PPI disruption in the dose-response format (Table 1, Figure 4C). Compound **24** contains a promiscuous maleimide, which is well known to react indiscriminately with proteins. Compound **4** had poor solubility (25 μM, data not shown), with both **4** and **32** displaying inconsistent GST-PD data (Figure 4B and Figure S2A–B). Therefore, these compounds were not progressed further.

**Figure 4.**
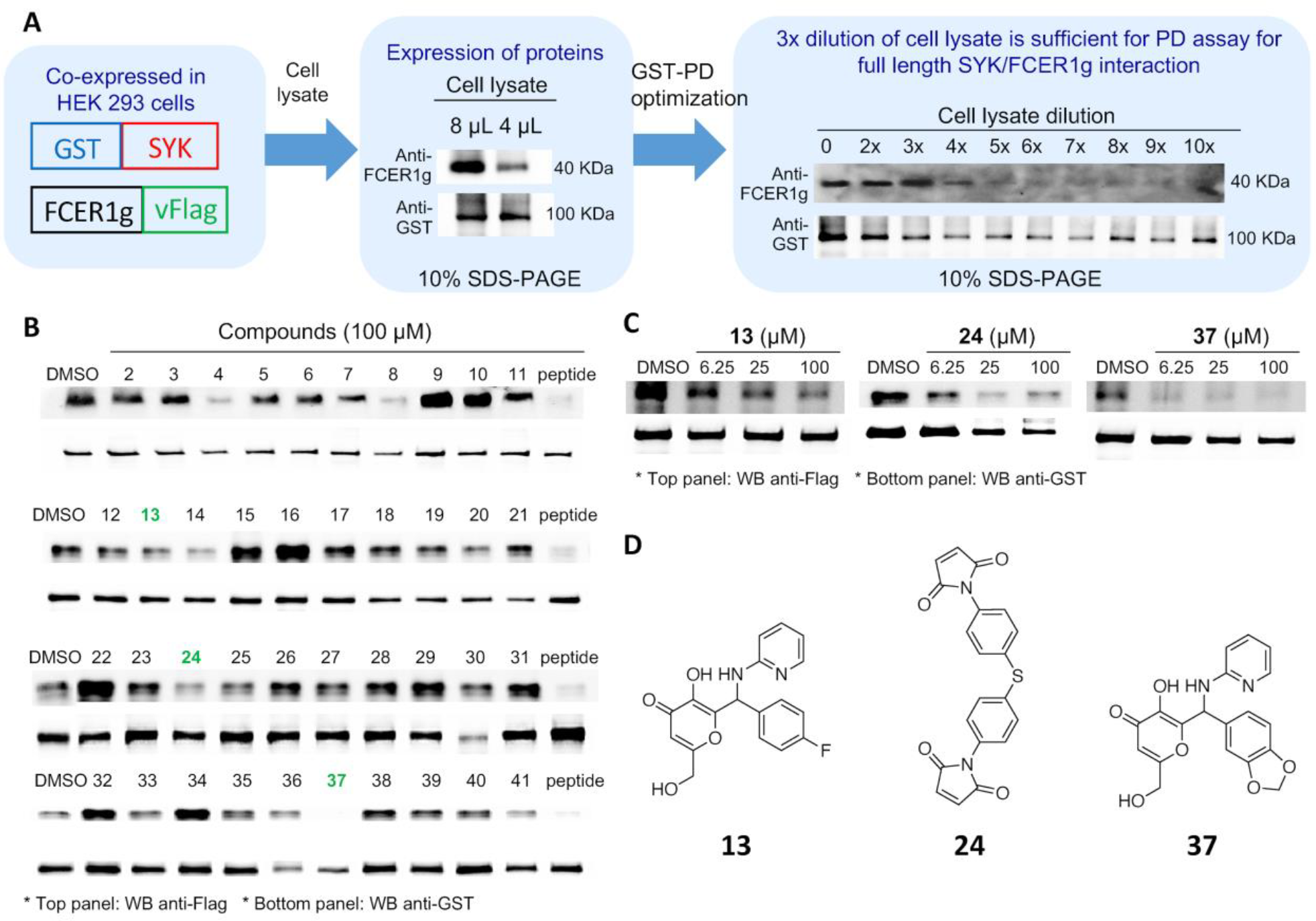
Identification of 3 compounds that inhibit the interaction of SYK and FCER1G in a GST-SYK and FCER1G-Flag pulldown assay. **(A)** Optimization of the GST-SYK and FCER1G-Flag pulldown assay in HEK293 cells. **(B)** Compounds **13, 24**, and **37** were able to inhibit the interaction between SYK-GST and FCER1G-Flag in a pulldown assay at a single concentration (100 μM). **(C)** Compounds **13, 24**, and **37** were tested via GST-pulldown in a dose-response fashion. **(D)** Chemical structures of compounds **13, 24**, and **37**.

**Table 1.**
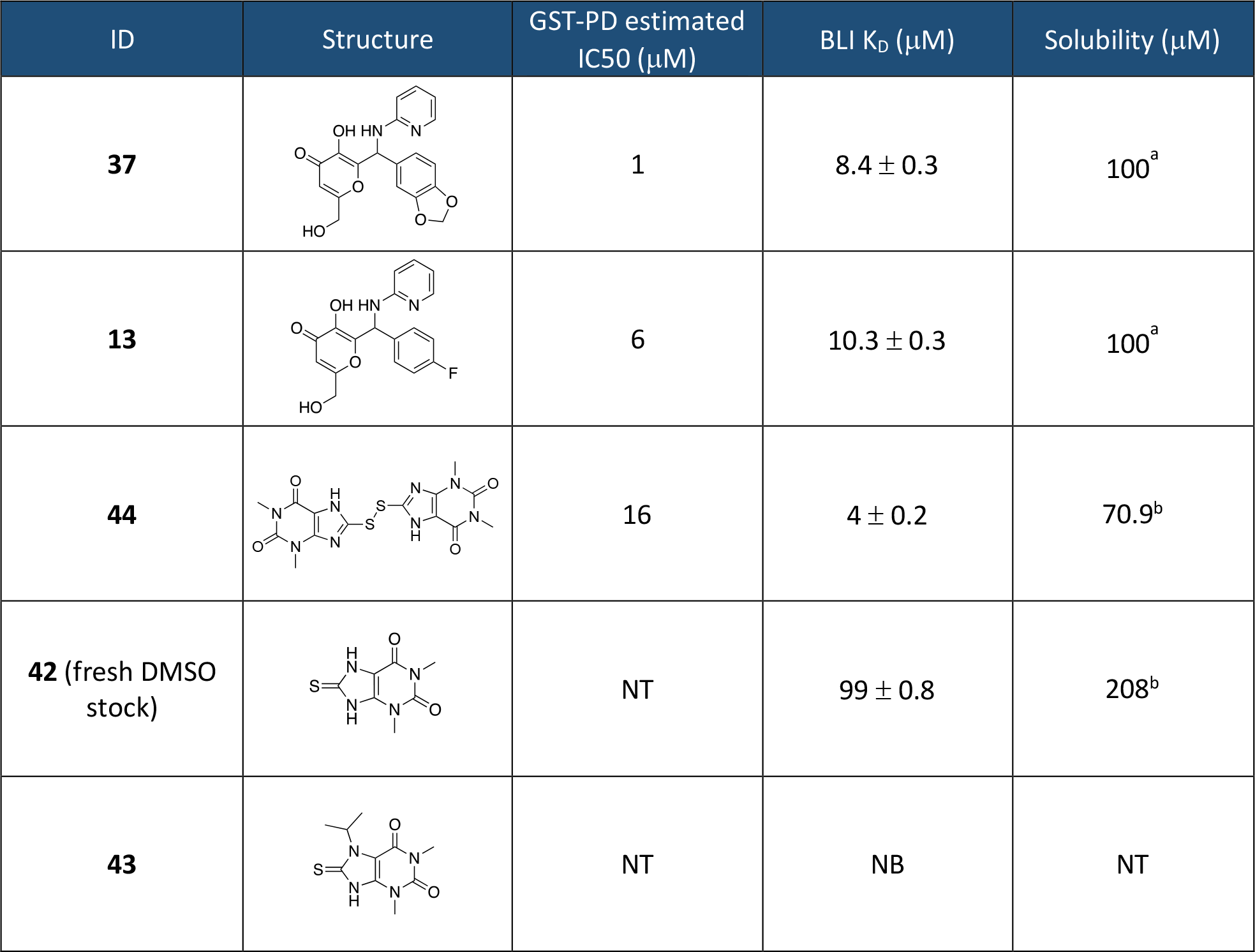
Biochemical and biophysical data for compounds **37, 13, 42, 43**, and **44**. The estimated IC_50_ for GST-PD is calculated via the western blot densitometry. ^a^ Solubility calculated with a nephelometer (Figure S5). ^b^ Kinetic solubility calculated by Analiza, Inc.

| ID | Structure | GST-PD estimated $IC_{50}$ (μM) | BLI $K_D$ (μM) | Solubility (μM) |
| --- | --- | --- | --- | --- |
| **37** | | 1 | $8.4 \pm 0.3$ | $100^a$ |
| **13** | | 6 | $10.3 \pm 0.3$ | $100^a$ |
| **44** | | 16 | $4 \pm 0.2$ | $70.9^b$ |
| **42** (fresh DMSO stock) | | NT | $99 \pm 0.8$ | $208^b$ |
| **43** |  | NT | NB | NT |

To validate direct binding of hit compounds **13, 32**, and **37** to SYK-tSH2, we performed an orthogonal biophysical protein-small molecule direct binding assay, BLI. A biotinylated version of SYK-tSH2 (SYK-tSH2-c022; residues M6-269) was used, which lacked the 6xHis tag (Figure S1C). Compounds **13** and **37** were shown to bind to SYK-tSH2 with K_d_ values of 8.4 μM and 10.3 μM, respectively (Table 1, Figure S3). In addition to the inconsistent GST-PD results previously mentioned, compound **32** did not bind to the SYK-tSH2 domains and therefore it was not progressed further.

### Compounds 13 and 37 covalently modify SYK-tSH2

The HTS hits **13** and **37** contain a kojic acid moiety, which is proposed to react covalently with proteins in the literature (38). As a result, we thought that it was likely these compounds are also reacting covalently with SYK-tSH2. Compounds **13** and **37** were incubated with SYK-tSH2 at 100 μM for 1 h at room temperature and a covalent reaction was revealed by mass spectrometry (Figure 5A). When bound to the protein, these kojic acid-derived compounds have a 94 Da decrease in mass due to the loss of the pyridin-2-amine leaving group to reveal a Michael acceptor that can react covalently with cysteine residues (Figure 5B). **13** and **37** formed between one and five and between two and seven mass adducts of 248 and 274 Da respectively, suggesting that these compounds react indiscriminately with cysteine residues and likely other nucleophilic amino acid residues due to the mass adducts observed totaling above the number of available cysteines (4).

**Figure 5.**
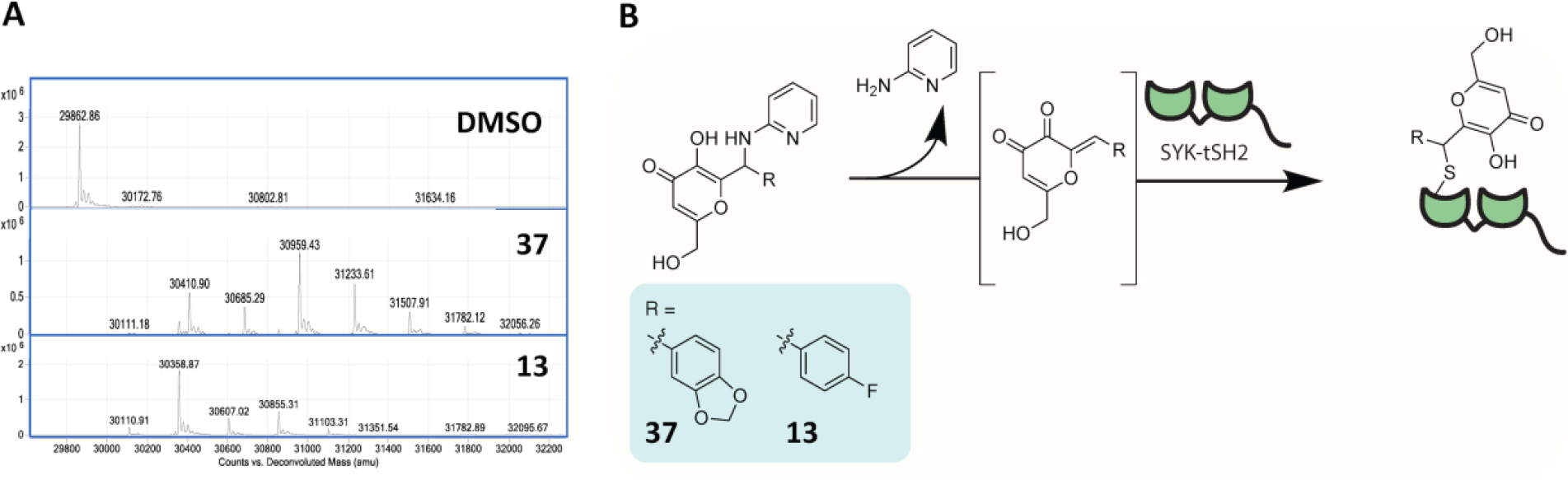
Compounds **13** and **37** react covalently with SYK-tSH2. **(A)** Mass spectrometry analysis of SYK incubated with compounds **37** and **13** at 100 μM for 1 h at room temperature. **(B)** Hypothesized mechanism of inhibition of **37** and **13** with SYK-tSH2.

### Hit expansion of 40 putative hits leads to identification of thiouric acid compounds that covalently modify SYK-tSH2 via oxidative disulfide bond formation

To expand on the initial 40 putative hits that were identified via uHTS (Figure 3D), additional commercially available structural analogues were ordered and screened via TR-FRET and differential scanning fluorimetry (DSF) (Table S1). Two analogues of primary hit **39** (IC_50_ = 27.7 μM) from uHTS, compounds **42** and **43** (Figure 6A–B), significantly destabilized SYK-tSH2 protein in the DSF assay and demonstrated single-digit micromolar IC_50_ values in the TR-FRET assay (Table S1). **43** is a close analogue of the primary hit **39**, varying only in methylation of the imide nitrogen versus **39**, and demonstrated a TR-FRET IC_50_ of 3.0 ± 1.7 μM and a DSF *ΔT*_*m*_ of -6.7 ± 0.4°C. Strikingly, compound **42**, which also contains a methylated imide nitrogen but lacks the isopropyl group on the thiourea nitrogen versus **39**, demonstrated a TR-FRET IC_50_ of 1.9 ± 0.20 μM and a DSF *ΔT*_*m*_ of -17 ± 0.3°C (Table S1). The observation of a substantial reduction in protein melting temperature motivated us to determine if the compounds interacted covalently with the SYK-tSH2 domains. Additionally, since the thiourea in these small molecules is reported to react via oxidation, the compounds could be capable of forming a covalent disulfide bond with cysteine residues on SYK (39). We used compound **1**, which is a known covalent modifier of SYK-tSH2, as a positive control in the mass spectrometry analysis (27). Compound **1** could not be used as a positive control in the TR-FRET due to assay interference with the compound quenching the TR-FRET donor signal. Compound **1** formed one or two × 222 Da adducts with the protein, suggesting that under these experimental conditions **1** reacts with either one or two cysteine residues covalently. Incubation of **42** and **43** at 100 μM for 1 h with SYK-tSH2 revealed that only compound **42** covalently modifies SYK-tSH2, but oddly only with older DMSO stock solutions (Figure 6C). Despite the similar structures, **43** did not demonstrate covalent attachment. Adducts of two (422 Da) more than the protein mass after **42** incubation suggests that this compound is binding at two cysteine residue sites on the protein surface (Figure 6C). Confoundingly, mass spectrometry analysis of SYK-tSH2 with a freshly prepared DMSO stock of **42**, however, did not result in the previously observed mass adduct (Figure 6C). Analytical characterization of freshly prepared solutions of **42** and **43** versus the older DMSO stocks that had been used in the preliminary mass spectrometry analyses revealed that the compounds were monomeric when freshly dissolved but had formed corresponding disulfide dimers in DMSO over time (Figure S4). The freshly prepared DMSO stock solution of **42** did not form a covalent adduct (Figure 6D). Literature reports support that this type of chemical reaction can occur in DMSO, forming a disulfide bond (40, 41). The proposed active disulfide compound, **44**, was synthesized from **42** by heating it in DMF at 100 °C and then purifying via aqueous filtration (Figure 6B) (42). **44** exhibits poor organic solubility and, accordingly, its melting point temperature was >300 °C (43, 44). The kinetic solubility of **44** was measured as 70.9 μM, which as expected is less soluble than the monomer **42** at 208 μM, however, is in a reasonable range of solubility (Table 1). Like we had originally observed for the older DMSO stock solution of **42** (Figure 6C), **44** forms two to four mass adducts (422 Da and 844 Da) more than the protein mass when incubated SYK-tSH2 for 1 h (Figure 6D). Since the freshly made DMSO stock solutions of **42** did not covalently modify SYK-tSH2, we propose that it forms disulfide **44** over time and this disulfide compound is responsible for the observed activity. Additionally, **44** was potent in the TR-FRET assay, demonstrating an IC_50_ = 0.27 ± 0.02 μM (Figure 6F). When **44** was analyzed by mass spectrometry in dose-response format, it exhibited 50% labeling of 10 μM SYK-tSH2 at 30 μM **44** (Figure 6E). Due to the mass adducts of two to four (422 Da and 844 Da), we propose that **44** is forming a disulfide bond with between two or four cysteines on SYK-tSH2 (Figure 6G).

**Figure 6.**
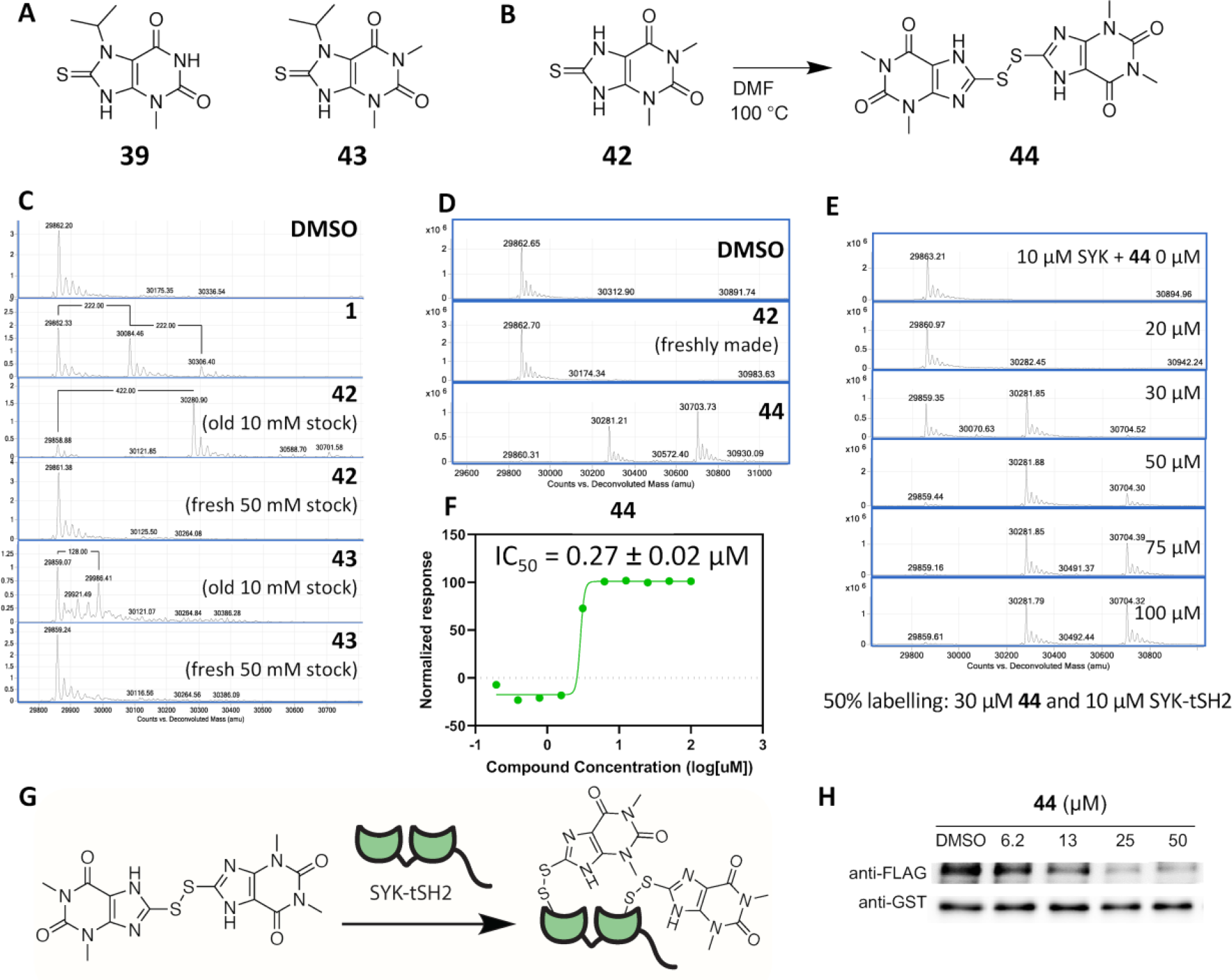
Compound **44** covalently modifies SYK-tSH2. **(A)** Chemical structures of initial HTS hit **39** and analogue **43. (B)** Synthesis of **44** from analogue **42. (C)** Mass spectrometry of SYK-SH2 incubated with covalent inhibitor **1** and old versus fresh stocks of **42** and **43** at 100 μM for 1 h at room temperature. Only the old DMSO stock of **42** reacts covalently with SYK-tSH2. **(D)** Mass spectrometry of SYK-tSH2 incubated with **42** and **44** at 100 μM for 1 h at room temperature. **(E)** Dose-response of **44** in covalently labelling SYK-tSH2, assessed by mass spectrometry. **(F)** Dose response curve of **44** in the TR-FRET assay (IC_50_ = 0.27 ± 0.02 μM). **(G)** Hypothesized mechanism of inhibition of **44** with SYK-tSH2. **(H)** Compound **44** disrupted the PPI via GST-pulldown in a dose-response fashion.

To confirm binding to SYK-tSH2 domains via an orthogonal assay the compounds **42** (as a fresh DMSO stock containing only the monomer), **43** and **44** were analyzed by BLI (Table 1, Figure S3). **44** demonstrated slightly more efficacious binding to SYK-tSH2 than the compounds identified from the HTS screen (**13** and **37**), with a K_d_ = 4.0 μM (Table 1). Compound **42** was also evaluated via BLI as a fresh DMSO stock, which should only contain the monomeric compound, and did not bind to the protein (K_d_ = 99 μM). Finally, consistent with the mass spectrometry analysis (Figure 6C), **43** did not bind to SYK-tSH2 when analyzed via BLI (Table 1). Additionally, compound **44** was able to disrupt the full-length interaction of SYK and FCER1G via a GST-PD assay (Table 1, Figure 6H).

### Compounds 13, 37, and 44 lack selectivity for SYK-tSH2

Mass spectrometry was used to determine the selectivity of the compounds identified as covalent inhibitors of SYK-tSH2 (**13, 37**, and **44**) against a select panel of proteins. Proteins MSN, TBXT, and SHIP1 were used for this experiment due to the knowledge that they contain accessible cysteines that have been targeted for prior covalent compound screening. The compounds were incubated at a concentration of 100 μM with the respective protein for 1 h at room temperature (Figure 7). As previously shown herein (Figure 5–6 and S6), compounds **13, 37**, and **44** form covalent mass adducts with SYK protein. **44** incubated with TBXT, which has two available cysteines, forms two mass adducts (420 Da). Compounds **13** and **37** both form two or three mass adducts, suggesting that the compound does not exclusively covalently react with cysteines on TBXT and reacts with additional residues. MSN, which has four available cysteines for modification, only forms two mass adducts (420 Da) with **44**. Intriguingly, compounds **13** and **37** only form one weak mass adduct with MSN (248 and 274 Da, respectively), with protein predominantly unlabeled by the two compounds. **44** was incubated at a lower concentration of 30 μM with SYK, TBXT, and MSN and resulted in the same formation of mass adducts as 100 μM incubation (Figure S6A, C, and D). Additionally, **44** was incubated with SHIP1, which has 10 available cysteine residues, and forms mass adducts of 8 or 10 (1678 or 2101 Da) (Figure S6B). Collectively these data indicates that **13, 37**, and **44** bind covalently to available cysteines on distinct proteins indiscriminately. Furthermore, compounds **13**, and **37** show mass adducts which are greater than the number of available cysteines, suggesting the potential for more promiscuous binding with amino acid residues.

**Figure 7.**
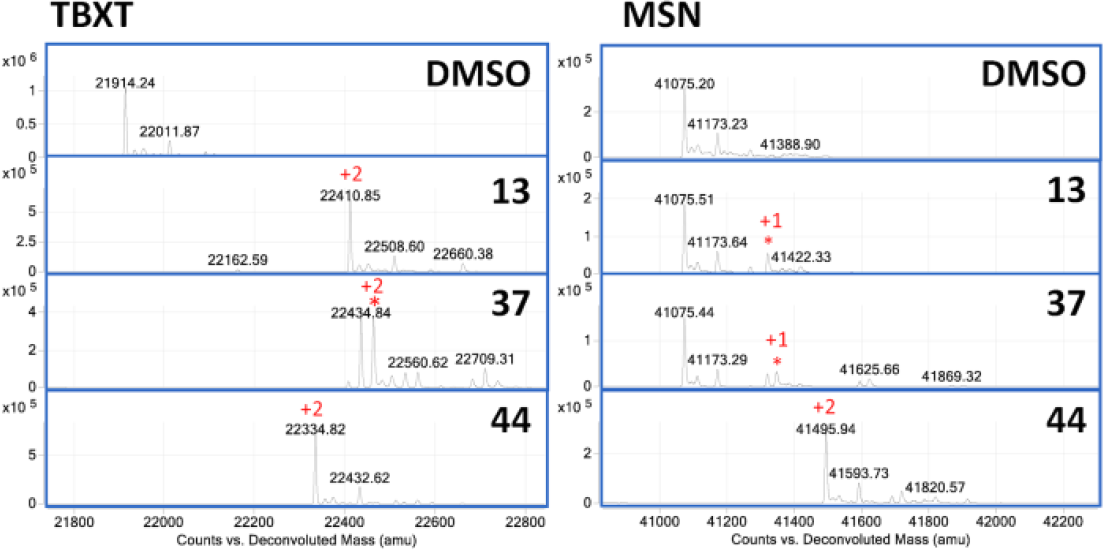
Compounds **13, 37**, and **44** (100 μM) bind covalently to TBXT and MSN via mass spectrometry. Number of mass adducts of the compound are indicated in red.

## Discussion

Herein, we describe the design and utilization of several biochemical and biophysical assays aimed at identification of inhibitors of the PPI between SYK-tSH2 and the FCER1G phospho-ITAM. As this interaction promotes activation of SYK and activated SYK has been implicated as a driver of AD pathology, pharmacological inhibition of this PPI is proposed to slow AD progression. Furthermore, SYK expression is elevated in AD-relevant regions of the brain in AD mouse models. We employed a biochemical TR-FRET assay as a high-throughput approach to identify putative PPI inhibitors. Since TR-FRET is an indirect method of detection, BLI, an orthogonal biophysical assay was enlisted to demonstrate direct binding of the inhibitors to SYK-tSH2. Mass spectrometry analysis confirmed that binding to SYK-tSH2 was covalent for some compounds (**13, 37**, and **44**), but not others (**42** and **43**). GST-PD assays confirmed that the PPI disruption was maintained when full length rather than truncated proteins were employed for these compounds.

We noted slight discrepancies in the data generated using the different assays. The TR-FRET assay identified many single-digit micromolar inhibitors of our PPI, including those compounds found in Table 1 and Table S1. However, not all of these were confirmed to bind to SYK-tSH2 via the BLI orthogonal biophysical assay and GST-PD assays. Most compounds pursued for further studies displayed at least single-digit micromolar potency in the TR-FRET assay. Generally, compounds that showed potent binding to SYK-tSH2 via BLI reacted covalently with the protein as assessed by mass spectrometry.

When structurally similar analogs of **39** were analyzed via DSF (Table S1), significant destabilization of SYK-tSH2 was observed with compounds **42** and **43**, and the most destabilizing compounds corresponded to the most potent TR-FRET value. Significant destabilization observed in the DSF assay motivated examination of a potential covalent mechanism of binding. Compound **42** (an old DMSO solution, mostly containing **44**) was shown to bind covalently to SYK-tSH2. However, compound **43** did not form a mass adduct corresponding to compound addition, despite the destabilizing DSF value. We found that compounds **42** and **43** form a disulfide in DMSO solution, which is responsible for the increased mass observed and covalent inhibition of SYK-tSH2. Evaluation of analogs of the initial HTS hit (**39**) led us to compound **44**, which was discovered to be the active component in older DMSO stocks of **42**. Monomeric **42** had formed a disulfide compound over time in DMSO, resulting in formation of **44. 44** has sub-micromolar potency in the TR-FRET assay, weak micromolar potency via BLI, and inhibited the GST-PD assay with weaker double-digit micromolar estimated IC_50_ values. The observed poor solubility of **44** compared to the monomeric **42** leads us to propose that the chemotype suffers from solubility issues (Table 1), which were magnified when moving into cellular lysate for GST-PD experiments. We propose that for all DMSO stocks containing the thiouric acid moiety found in **42** and **43** a percentage of the compound present will have formed the disulfide compound. In support of this reactivity, **42** is often used as a reactant in the synthesis of more elaborate compounds due to the oxidative nature of its sulfur atom (39, 45). We propose that the isopropyl group on **43** reduces its ability to form the active disulfide species as efficiently as **42** and/or that it exists in a dynamic equilibrium with its monomeric counterpart. A structurally similar analog of **42**, 8-thiouric acid, was co-crystallized with urate oxidase (UOX, PDB code: 3LBG) and demonstrated both non-covalent binding of the monomeric compound and a covalent disulfide bond formed between the monomeric compound and Cys35 (46). It is possible that the covalent modification of UOX was executed by a dimeric form of 8-thiouric acid, similar to what we observed for **42**, forming **44** and then labeling SYK-tSH2 (Figure 6G). Despite the covalent nature of these inhibitors, promisingly we did not observe indiscriminate binding to the cysteines on SYK-tSH2, with only two to four mass adducts observed up to a concentration of 100 μM. Compounds **13** and **37** that were identified from the primary HTS share the same chemotype, which contains a kojic acid derived core scaffold. **13** and **37** consistently exhibited single-digit micromolar values in TR-FRET, BLI, and GST-PD assays. The chemotype for these compounds is different to compound **44** and compound **1**, which bind via an oxidative disulfide and covalent aromatic substitution mechanism, respectively. Inhibitors containing the kojic acid core modify an unrelated protein - transcriptional enhanced associate domain (TEAD) (38). The reported mechanism that these compounds react with TEAD involves an initial retro Mannich reaction, revealing a Michael acceptor that can react covalently with an available cysteine residue on the protein (Figure 5B). Compounds **13** and **37** bind to cysteine(s) on SYK-tSH2, resulting in potent inhibition of our PPI. Multiple adducts are observed with these compounds, further suggesting that the compounds will likely bind promiscuously to readily available cysteine residues. Early identification of this indiscriminate labeling of another protein diminished our enthusiasm for these hits.

While **13, 37**, and **44** are confirmed to inhibit the PPI between SYK-tSH2 and the FCER1G peptide with micromolar potency, the selectivity of these small molecules are poor via mass spectrometry. Despite **44** binding to fewer amino acid residues on SYK-tSH2 than **13** and **37** observed by mass spectrometry, against a select panel of proteins with known available cysteines, no selectivity was observed. Despite the success inhibiting the PPI between SYK and FCER1G peptide, it is clear that more work needs to be pursued to improve the selectivity of these compounds. Furthermore, we were unable to detect an effect on SYK phosphorylation in a cellular inhibition assay (data not shown), and therefore these compounds would also need to be further optimized to improve potency and ensure cell penetrance. These compounds require further optimization to become good tools for interrogating the function of SYK and FCER1G in a cellular context and should only be used with caution.

## Conclusion

We have identified three confirmed (**13, 37**, and **44**) covalent inhibitors of SYK-tSH2 domain that disrupt its interaction with the FCER1G peptide when used at micromolar concentrations. These compounds bind to SYK-tSH2 when evaluated using biophysical methods and mass spectrometry. Additionally, they inhibit the interaction between full length SYK and FCER1G. These compounds highlight that covalent inhibition of SYK-tSH2 is a viable strategy to assess the implications of inhibiting the PPI between SYK and FCER1G. Intriguingly, we have identified covalent inhibition of SYK-tSH2 via two distinct mechanisms to that of compound **1**, which reacts via a covalent aromatic substitution reaction with cysteine. Cysteine reacts via a Michael addition on the alkene revealed on compounds **13** and **37**. The compounds undergo an initial retro Mannich reaction that generates the alkene, which is the reactive Michael acceptor species for covalent inhibition. With compound **44**, which is a disulfide compound, an oxidative reaction occurs forming a disulfide with an available cysteine residue. We have highlighted some limitations related to the reactivity of the kojic acid core shared by **13** and **37** and the disulfide containing compound **44** with proteins unrelated to SYK. In addition, the poor solubility of **44** in both aqueous and organic solutions as well as poor selectivity for the SYK-FCER1G interface limits its utility. Overall, we have demonstrated that the interaction between these two proteins can be disrupted via small covalent molecules that utilize distinct covalent mechanisms, identifying this strategy as a potential alternative approach towards the development of small molecule regulators of SYK signaling.

## Materials and Methods

### Plasmid constructs and peptides

Expression plasmids used in protein purification were generated by PCR amplification of the tandem SH2 domains of human SYK (residues M6-N269) from the Mammalian Gene Collection cDNA library (IMAGE:3870426). The PCR product was cloned into the *E. coli* expression vector pNIC28-Bsa4 or pNIC-Bio3 using ligation independent cloning. Cloning into both vectors gives rise to an N-terminal fusion of SYK with a 6xHis tag and a TEV protease cleavage site (SYKA-c020), while the pNIC-Bio3 construct contains in addition a C-terminal AviTag fusion. TR-FRET assays were performed using an FITC-conjugated peptide containing the FCER1G ITAM sequence with two phosphotyrosines (FCER1G-phospho-ITAM[62-81]-FITC: DGV(pY)TGLSTRNQET(pY)ETLKH-FITC). A non-phosphorylated ITAM sequence was used as a control (FCER1G-nonphospho-ITAM[62-81]-FITC: DGVYTGLSTRNQETYETLKH-FITC). Biophysical assays employed a phosphorylated ITAM without FITC conjugation (FCER1G-phospho-ITAM[62-81]: DGV(pY)TGLSTRNQET(pY)ETLKH). Peptides were custom synthesized by LifeTein. GST-, VF-tagged human SYK and FCER1G plasmids for mammalian expression were generated using Gateway cloning technology (Invitrogen) as described previously (47). The DNA was purified using ZymoPURE Plasmid Maxiprep Kit (D4203, Zymo Research).

### iPSC data

iPSCs were cultured and differentiated to neurons and microglia as previously described (48, 49). For neuronal differentiation, iPSCs were plated on Matrigel coated 6-well plates at a density of 3.5 million cells per well and fed with Basal Neural Maintenance Media (1:1 DMEM/F12 + glutamine media/neurobasal media, 0.5% N2 supplement, 1% B27 supplement, 0.5% GlutaMax, 0.5% insulin-transferrin-selenium, 0.5% NEAA, 0.2% β-mercaptoethanol; Gibco) + 10mM SB-431542 + 0.5mM LDN-193189 (Biogems). Cells were fed daily for seven days. On day eight, cells were incubated with Versene, gently dissociated using cell scrapers, and passaged at a ratio of 1:3. On day nine, media was switched to Basal Neural Maintenance Media and fed daily. On day 13, media was switched to Basal Neural Maintenance Media with 20 ng/mL FGF (R&D Systems) and fed daily. On day sixteen, cells were passaged again at a ratio of 1:3. Cells were fed until approximately day 23. At this time, cells were FACS sorted to obtain the CD184/CD24 positive, CD44/CD271 negative neural precursor cell (NPC) population. Following sorting, NPCs were expanded for neural differentiation. For cortical neuronal differentiation, NPCs were plated out in 10cm cell culture dishes at a density of 6 million cells/10 cm plate. After 24 h, cells were switched to Neural Differentiation media (DMEM-F12 + glutamine, 0.5% N2 supplement, 1% B27 supplement, 0.5% GlutaMax) + 0.02 μg/mL brain-derived neurotrophic factor (PeproTech) + 0.02 μg/mL glial-cell-derived neurotrophic factor (PeproTech) + 0.5 mM dbcAMP (Sigma Aldrich). Media was refreshed twice a week for three weeks. After three weeks, neurons were selected for CD184/CD44/CD271 negative population by MACS sorting and plated for experiments. For microglial differentiation iPSCs were differentiated into hematopoietic progenitor cells (HPCs) using the STEMdiff™ Hematopoietic Kit (05310, Stem cell Technologies). HPCs were either frozen using Bambanker HRM freezing media (BBH01, Bulldog Bio) or further differentiated into MGLs. HPCs were cultured in microglia differentiation medium comprised of DMEM/F12 (11039047, Thermo Fisher Scientific), B27 (17504-044, Thermo Fisher Scientific), N2 (17502-048, Thermo Fisher Scientific), insulin-transferrin-selenite (41400045, Thermo Fisher Scientific), non-essential amino acids (#11140050; Thermo Fisher Scientific), Glutamax (#35050061; Thermo Fisher Scientific), human insulin (I2643-25mg, Sigma) and monothioglycerol (M1753, Sigma) supplemented with 25 ng/ml human M-CSF (PHC9501, Thermo Fisher Scientific), 50 ng/ml TGF-β1 (130-108-969, Miltenyl), and 100 ng/ml IL-34 (200-34, Peprotech). After 24 days in this medium, two additional cytokines, 100 ng/ml CD200 (C311, NovoProtein) and 100 ng/ml CX3CL1 (300-31, PeproTech), were added to the medium described above to mature MGLs. MGLs were cultured in this new medium for an additional week and then harvested for experiments.

For mRNA expression analysis, RNA was extracted using the Trizol (15596018, Thermo Fisher Scientific) and first strand cDNA synthesis was performed using the iScript cDNA Synthesis kit (1708890, Bio-Rad). Quantitative PCR (qPCR) was performed with SYBR Green Master Mix (A46012, Thermo Fisher Scientific). SYK and FCER1G values were normalized to the geometric mean of the housekeeping genes, RPL13 and CYC1. qRT-PCR results were calculated using the 2^-DD*CT*^ method and presented as fold change from DMSO following the procedure described by Rao and co-workers (50).

Primer sequences:

SYK Primer #4:

Forward 5’GAGAAAGGAGAGCGGATGGG 3’

Reverse 5’ GGGCCTGTTTTCCACATCGT 3’

SYK Primer #8

Forward 5’ AGGTTTCCATGGGCATGAAGT 3’

Reverse 5’ CATGGGTCTGGGCCTTGTAG 3’

FCER1G Primer #1

Forward 5’ TGGTGTTTACACGGGCCTGA 3’

Reverse 5’ CCATGAGGGCTGGAAGAACC 3’

FCER1G Primer #2

Forward 5’ CGATCTCCAGCCCAAGATGA 3’

Reverse 5’ CCTTTCGCACTTGGATCTTCA 3’

### Bioinformatics

Risk score calculation and biological domain definitions are described in Cary *et al* (33). Pre-computed Target Risk Scores (syn25575156, v13), genetic risk scores (syn26844312, v6), and multi-omics risk scores (syn22758536, v9), were downloaded from Synapse. Single cell data (SEAAD_MTG_RNAseq_final-nuclei.2022-08-18.h5ad) were downloaded from the provided Amazon Web Services S3 bucket (s3://sea-ad-single-cell-profiling/MTG/RNAseq/) and processed with a custom script. Briefly, the script groups single cells based on the Supertype label as well as the Cognitive Status of the donor and computes an average expression for each gene as well as the fraction of cells with > 0 counts for that gene. Results were plotted using custom R scripts. All code used to process these data are available (github.com/caryga/TREATAD_target_reports).

### Protein purification

SYK-tSH2 protein was produced using *E. coli* BL21(DE3)-R3-pRARE cells grown in terrific broth. Prior to harvesting, cells were shifted to 18°C for 16 h after IPTG induction. For biotinylation of SYKA-c022 containing the AviTag, biotin (300 μM final) was added to the culture at the same time as IPTG addition. Cells were lysed by sonication in Lysis Buffer containing 50 mM HEPES (pH 7.5), 500 mM NaCl, 10 mM imidazole, 5% glycerol, 1 mM TCEP. SYK-tSH2 protein was bound to equilibrated Ni-IDA resin for 1 h before two batch washes in Lysis Buffer. Beads were subsequently loaded onto a drip column, followed by washing with Lysis Buffer containing 30 mM imidazole, before finally eluting bound protein with Lysis Buffer containing 300 mM imidazole. For TR-FRET assays, the 6xHis tag on SYKA-c020 was left intact. For biophysical assays, the 6xHis tag on SYKA-c020 or SYKA-c022 was cleaved off by the addition of a 1:10 mass ratio of 6xHis-TEV protease while undergoing dialysis (10k MWCO) in Lysis Buffer lacking imidazole for 16 h at 4°C. TEV protease and the cleaved 6xHis tag was subsequently removed with Ni-IDA resin equilibrated in Lysis Buffer. The molecular mass of purified proteins was confirmed by mass spectroscopy.

### Kinetic Solubility

Analiza, Inc. analyzed the kinetic solubility of a 5 mM DMSO stock of **44** and a freshly prepared 10 mM stock of **42** dissolved in phosphate buffered saline (PBS) solution at pH 7.4. Following 24 h incubation in a Millipore solubility filter plate, samples were vacuum filtered, and the filtrates collected for analysis. Filtrates were injected into the nitrogen detector for quantification via total chemiluminescent nitrogen determination (CLND). Filtrates were quantified with respect to a calibration curve generated using standards that span the dynamic range of the instrument. The reported solubility value has been corrected for background nitrogen present in the media and DMSO.

### TR-FRET Assay

TR-FRET experiments were performed in 20 μL reactions, using 384-well shallow-well microplates (PerkinElmer). Final reaction components contained 6xHis-SYK-tSH2 (SYKA-c020 tagged) (4 nM), FITC-conjugated FCER1G peptide (8 nM), Tb-conjugated anti-6xHis antibody (53 ng/ml; Cisbio) in assay buffer containing 25 mM HEPES (pH 7.5), 200 mM NaCl, 0.1% BSA and 0.05% Tween-20. After a 2h incubation (at room temperature), TR-FRET signals were measured on a BMG Labtech PHERAstar FSX reader using a Lanthascreen Optics Module. A 200 μs delay was used after excitation with a flash lamp before measurement of fluorescence emission at 490 and 520 nM. TR-FRET ratios of fluorescent intensity at 520 nm to 490 nm were calculated. The half maximal inhibitory concentration (IC50) was determined by fitting a four parametric logistic curve to the data.

### High-throughput TR-FRET Assay

The TR-FRET assay for 6xHis-SYK-tSH2 (SYKA-c020 tagged) and FITC-FCER1G peptide interaction was miniaturized and optimized into a 1536-well format for uHTS (details of the assay miniaturization will be described in a separate publication). The uHTS was performed in black 1536-well plate (3724, Corning costar) with a total volume of 5 μL in each well. 5 μL of the reaction mixture containing optimized concentrations of protein and peptide (6xHis-SYK-tSH2 protein: 2 nM, FITC-p-FCER1G: 6 nM, and anti-His-Tb 1:1000) was dispensed into black 1536-well plates using multiple-drop Combi dispenser (5840320, Thermo). The FITC-p-FCER1G and anti-His-Tb without His-SYK protein was used as background control. 0.1 μL of library compound dissolved in DMSO was added using pintool integrated with Beckman NX (Beckman Coulter, Brea, CA). The final compound concentration was 20 μM and the final DMSO concentration was 2%. The TR-FRET signals were measured using PHERAstar FSX plate reader (BMG) as described above. A total of 138,214 compounds from two chemical diversity libraries (ChemDiv and Asinex) at Emory Chemical Biology Discovery Center (ECBDC) have been screened.

Screening data were analyzed using Bioassay software from CambridgeSoft (Cambridge, MA). The S/B and Z’ in uHTS format were calculated for each screening plate. The effect of compound on the interaction TR-FRET signal was expressed as % of Control and calculated as the following equation:

% of Control = (F compound-F Background)/(F control – F background) × 100

Where F control and F background are the average TR-FRET signal from highest signal (His-SYK protein and FITC-p-FCER1G) and background without His-SYK protein (lowest signal), respectively. F compound is the TR-FRET signal for His-SYK/FITC-p-FCER1G interaction in the presence of library compound. Compounds that caused % Control < 60 were defined as primary positives.

### Differential Scanning Fluorimetry (DSF)

SYK-tSH2 protein (SYKA-c020 cleaved), lacking the N-terminal 6xHis tag, was diluted in assay buffer (2 μM in 10 mM HEPES, 150 mM NaCl) containing Sypro Orange (1 in 1000 dilution; Thermofisher). Compounds, control peptides (1 μM), or DMSO was added to SYK-tSH2 to 2% of final volume and the mixture was incubated for 30 min on ice. Melting curves were obtained on an Mx3005p qPCR machine (Agilent), ramping up from 25 to 95 °C, at 1°C min^−1^. Data were fitted GraphPad Prism to the Boltzmann equation. The *T*_*m*_ was calculated by determining the maximum value of the first derivative of fluorescence transition.

### GST-PD

The dose response confirmatory GST pulldown with test compounds was performed using the following reagents: Glutathione Sepharose 4B (17075605, Cytiva Sweden AB); WB primary antibodies: GST-Taq (2.6H1) Mouse mAb (2624S, Cell Signaling) 1:1000 dilution in TBST; Monoclonal Anti-FLAG M2-HRP antibody (A8592-1MG, Sigma) 1:1000 dilution in TBST; FcεR1γ mouse monoclonal antibody (SC-390222, Santa Cruz). WB secondary antibodies: peroxidase-conjugated AffiniPure Goat Anti-Mouse IgG (H+L) (115-035-003, Jackson Immuno Research) 1:5000 dilution in TBST. GST pulldown was performed as described with minor changes (51). HEK293T (CRL-3216, ATCC) were cultured in Dulbecco’s modified Eagle’s medium with 4.5 g/L glucose, L-glutamine, and sodium pyruvate (10–013-CV, Corning) supplemented with 10% fetal bovine serum and 1% penicillin/streptomycin solution (30–002-CI, CellGro). Cells were incubated at 37 °C in humidified conditions with 5% CO_2_. The HEK293T cells were transfected using 1 mg/mL Polyethylenimine (PEI; 23966, Polysciences) in a ratio of 3 μL to 1 μg DNA. The proteins were expressed for 48 h at 37 °C. Cell pellets were re-suspended in the SYK lysis buffer (25 mM HEPES, pH 7.5, 200 mM NaCl, 0.5% Triton X100 (T9284-1L, Sigma-Aldrich), followed by sonication for 10 s (5 s on, 5 s off) at 4 °C. Cells were pelleted by 10 min centrifugation at 13,000 rpm at 4 °C. The cleared lysates were first pre-incubated with compounds for 30 min and then incubated with glutathione sepharose 4B beads (17075605, Cytiva Sweden AB) at 4 °C for 2 h with rotation. After the incubation, beads were washed three times with the SYK lysis buffer, eluted by boiling in sodium dodecyl sulfate-polyacrylamide gel electrophoresis (SDS–PAGE) loading buffer (1610737, Bio-Rad), and analyzed by western blotting.

### Bio-Layer Interferometry

Bio-Layer Interferometry (BLI) for compound/protein direct binding was performed using the Octet RED384 system (Sartorius BioAnalytical Instruments) and biotinylated SYK-tSH2 protein (SYKA-c022 cleaved/biotinylated) (as previously described). The biotinylated SYK-tSH2 (50 μg/mL) was loaded onto super streptavidin sensors in 50 μL loading buffer (PBS + 0.02% Tween-20) for 10 min resulting in a 14 nm loading density. The assay was performed as described in (52).

### Compound solubility with Nephelometer

Stock compound (10 mM in DMSO) was serial diluted in PBS (21-040-CV, Corning) in 96-well Costar Assay plate (3904, Corning). After incubating compound in PBS for 90 min at room temperature, the Nephelometric turbidity units (NTUs) were measured with NEPHELOstar (BMG LABTECH, Cary, North Carolina). 0.2 s was used for a measure time and PBS only as the background. Each sample was tested in triplicate. Dose-response data was analyzed by Graphpad and the solubility concentration is defined at the concentration that increases the NTUs at the turning point of the curve.

### Covalent labelling and mass spectrometry of proteins

Proteins were diluted to 80 μg/ml in buffer containing 50 mM HEPES (pH 7.5) and 200 mM NaCl before the addition of compounds to test for covalently labelling, giving a final concentration of 2% DMSO in the reaction. After incubation for 1 h at room temperature, the reaction was stopped by the addition of 4 volumes of 0.2% formic acid. Reversed-phase chromatography was performed in-line prior to mass spectrometry using an Agilent 1100 HPLC system (Agilent Technologies inc. – Palo Alto, CA, USA). Protein samples in formic acid were injected (50 μL) on to a 2.1 mm x 12.5 mm *Zorbax* 5um 300SB-C3 guard column housed in a column oven set at 40 °C. The solvent system used consisted of 0.1% formic acid in ultra-high purity water (Millipore) (solvent A) and 0.1 % formic acid in methanol (LC-MS grade, Chromasolve) (solvent B). Chromatography was performed as follows: Initial conditions were 90 % A and 10 % B and a flow rate of 1.0 ml/min. After 15 s at 10 % B, a two-stage linear gradient from 10 % B to 80 % B was applied, over 45 s and then from 80% B to 95% B over 3 s. Elution then proceeded isocratically at 95 % B for 1 min 12 s followed by equilibration at initial conditions for a further 45 s. Protein intact mass was determined using a 1969 MSD-ToF electrospray ionisation orthogonal time-of-flight mass spectrometer (Agilent Technologies Inc. – Palo Alto, CA, USA). The instrument was configured with the standard ESI source and operated in positive ion mode. The ion source was operated with the capillary voltage at 4000 V, nebulizer pressure at 60 psig, drying gas at 350°C and drying gas flow rate at 12 L/min. The instrument ion optic voltages were as follows: fragmentor 250 V, skimmer 60 V and octopole RF 250 V.

### Purchase of commercial compounds

Reagents were obtained from verified commercial suppliers and used without further characterization or purification. **1** (Compound B in Visperas *et al* (27), CAS:378201-55-9). **37** (PubChem ID: 24319153) was purchased from Vitas-M Laboratory (Vendor ID: STL228089). **13** (PubChem ID: 24337874) was purchased from Enamine (Vendor ID: Z56176033). Compounds **2** – **41** were purchased from ChemDiv, Chembridge, Vitascreen, LifeChemicals, and Enamine. Other UNC compounds (**42, 43, 46**–**60**) were purchased from ChemSpace.

### Chemistry

#### General Information

Temperatures are reported in degree Celsius (°C); the solvent was removed via a rotary evaporator under reduced pressure; and thin layer chromatography was used to monitor the progress of reactions that were executed unless otherwise noted. The following abbreviations are used in schemes and/or experimental procedures: μmol (micromoles), mg (milligrams), equiv (equivalent(s)), and h (hours). ^1^H NMR and/or additional microanalytical data were collected for intermediates and final compounds to confirm their identity and assess their purity. ^1^H spectra were obtained in DMSO-*d*_6_ and recorded using Varian or Bruker spectrometers. Magnet strength is indicated in the line listing. Peak positions are listed in parts per million (ppm) and calibrated versus the shift of the indicated deuterated solvent; coupling constants (*J* values) are reported in hertz (Hz); and multiplicities are included as follows: singlet (s). Purity was determined using high-performance liquid chromatography (HPLC). All final compounds are >95% pure by HPLC analysis.

#### Synthesis of 44

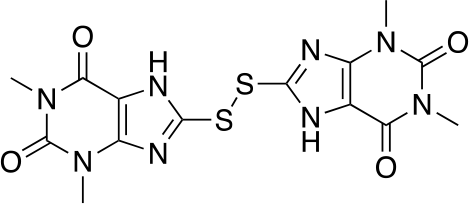

To a flask was added 1,3-dimethyl-8-thioxo-3,7,8,9-tetrahydro-1*H*-purine-2,6-dione (100 mg, 1 equiv., 471 μmol) and DMF (2.36 mL) and the reaction refluxed at 100 ° C for 4 h until a yellow precipitate crashed out. The precipitate was washed with H_2_O under vacuum filtration and further dried under vacuum to yield the desired product as a yellow solid 8,8’-disulfanediylbis(1,3-dimethyl-3,7-dihydro-1*H*-purine-2,6-dione) (38 mg, 19 %). ^1^H NMR (400 MHz, DMSO-*d*_6_): δ 3.42 (s, 3H), 3.25 (s, 3H). LCMS Calculated for C_14_H_15_N_8_O_4_S_2_ [M+H]^+^: 423.06. Found: 423.0. MP: >300 °C.

## Acknowledgements

The authors thank Drs. Allan I. Levey, Lara M. Mangravite, Haian Fu, Opher Gileadi, Aled Edwards, Frank Longo, Gregory W. Carter, Tim M. Willson, and Stephen V. Frye for insightful discussions and suggestions during the development of these resources. The Target Enablement to Accelerate Therapy Development for Alzheimer’s Disease (TREAT-AD) Consortium was established by the National Institute on Aging (NIA). The research reported in this manuscript was led by the Emory-Sage-SGC TREAT-AD center and supported by grant U54AG065187 from the NIA. The Structural Genomics Consortium is a registered charity (number 1097737) that receives funds from AbbVie, Bayer Pharma AG, Boehringer Ingelheim, Canada Foundation for Innovation, Eshelman Institute for Innovation, Genome Canada, Genentech, Innovative Medicines Initiative (EU/EFPIA), Janssen, Merck KGaA Darmstadt Germany, MSD, Novartis Pharma AG, Ontario Ministry of Economic Development and Innovation, Pfizer, São Paulo Research Foundation-FAPESP, Takeda, and Wellcome. The funders did not play a role in study design, data collection and interpretation, decision to publish, or manuscript preparation.

## Members of Emory-Sage-SGC TREAT-AD Center

Ishita Ajith^1^, Joel K. Annor-Gyamfi^2^, Jeff Aube^2^, Alison D. Axtman^2^, Frances M. Bashore^2^, Ranjita S. Betarbet^3^, Juan Botas^4^, William J. Bradshaw^1^, Paul E. Brennan^1^, Peter J. Brown^2^, Robert R. Butler 3rd^5^, Jacob L. Capener^2^, Gregory W. Carter^6^, Gregory A. Cary^6^, Catherine Chen^4^, Rachel Commander^3^, Sabrina Daglish^2^, Suzanne Doolen^7^, Yuhong Du^3^, Aled M. Edwards^8^, Michelle E. Etoundi^4^, Kevin J. Frankowski^2^, Stephen V. Frye^2^, Haian Fu^3^, Opher Gileadi^1^, Marta Glavatshikh^2^, Jake Gockley^9^, Katerina Gospodinova^1^, Anna K. Greenwood^9^, Peter A. Greer^10^, Lea T. Grinberg^11^, Shiva Guduru^2^, Levon Halabelian^8^, Crystal Han^5^, Brian Hardy^2^, Laura M. Heath^9^, Stephanie Howell^2^, Andrey A. Ivanov^3^, Suman Jayadev^12^, Vittorio L. Katis^1^, Stephen Keegan^6^, May Khanna^13^, Dmitri Kireev^2^, Carl LaFlamme^14^, Karina Leal^9^, Tom V. Lee^4^, Tina M. Leisner^2^, Allan I. Levey^3^, Qianjin Li^3^, David Li-Kroeger^4^, Zhandong Liu^4^, Benjamin A. Logsdon^9^, Frank M. Longo^5^, Lara M. Mangravite^9^, Peter S. McPherson^14^, Richard M. Nwakamma^3^, Felix O. Nwogbo^2^, Carolyn A. Paisie^6^, Arti Parihar^12^, Kenneth H. Pearce^2^, Kun Qian^3^, Min Qui^3^, Stacey J Sukoff Rizzo^7^, Karolina A. Rygiel^1^, Julie Schumacher^5^, David D. Scott^15^, Nicholas T. Seyfried^3^, Joshua M. Shulman^4^, Ben Siciliano^3^, Arunima Sikdar^2^, Nathaniel Smith^4^, Michael Stashko^2^, Judith A. Tello Vega^15^, Dilipkumar Uredi^2^, Dongxue Wang^3^, Jianjun Wang^3^, Xiaodong Wang^2^, Zhexing Wen^3^, Jesse C. Wiley^9^, Alexander Wilkes^1^, Charles A. Williams^12^, Timothy M. Willson^2^, Aliza Wingo^3^, Thomas S. Wingo^3^, Novak Yang^3^, Jessica E. Young^12^, Miao Yu^6^, Elizabeth L. Zoeller^3^

^1^University of Oxford, Oxford, OX3 7FZ, UK

^2^University of North Carolina, Chapel Hill, NC 27599, USA

^3^Emory University School of Medicine, Atlanta, GA 30322, USA

^4^Baylor College of Medicine, Houston, TX 77030, USA

^5^Stanford University School of Medicine, Stanford, CA, 94305, USA

^6^The Jackson Laboratory, Bar Harbor, ME 04609, USA

^7^University of Pittsburgh School of Medicine, Pittsburgh, PA 15219, USA

^8^University of Toronto, Toronto, ON M5G 1L7, Canada

^9^Sage Bionetworks, Seattle, WA, 98121, USA

^10^Queen’s University, Kingston, Ontario, ON K7L 3N6, Canada

^11^University of California, San Francisco, San Francisco, CA 94143, USA

^12^University of Washington, Seattle, WA 98109, USA

^13^New York University, New York, NY 10010, NY, USA

^14^McGill University, Montreal, QC H3A 2B4, Canada

^15^University of Arizona, Tucson, AZ 85724, USA

