## Supplementary Information for "Characterization of covalent inhibitors that disrupt the interaction between the tandem SH2 domains of SYK and FCER1G phospho-ITAM"

### Supplemental Figures and Tables

| UNC# | Structure | TR-FRET ($IC_{50}$) µM | DSF ($\Delta T_m$) |
|---|---|---|---|
| **39** | 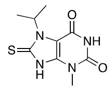 | 27.17[a] | NT |
| **43** | 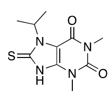 | 1.9 ± 0.20 | -16.6 ± 0.3 |
| **42** | 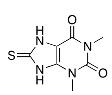 | 3.0 ± 1.7 | -6.7 ± 0.4 |
| **45** | 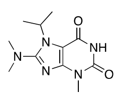 | > 100 | NT |
| **46** | 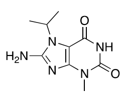 | > 100 | NT |
| **47** | 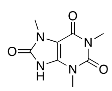 | >100 | NT |
| **48** | 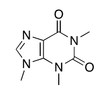 | >100 | NT |
| **49** | 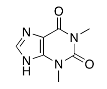 | >100 | NT |
| **50** | 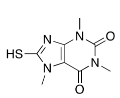 | 3.57 ± 0.20 | NT |
| **51** | 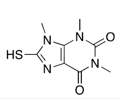 | 2.93 ± 0.16 | NT |

| UNC# | Structure | TR-FRET ($IC_{50}$) µM | DSF ($\Delta T_m$) |
|---|---|---|---|
| **52** | 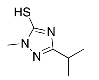 | 5.41± 0.06 | NT |
| **53** | 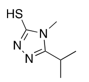 | 4.94± 0.23 | NT |
| **54** | 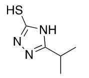 | 12.52± 0.57 | NT |
| **55** | 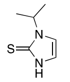 | 9.7 ± 4.3 | NT |
| **56** | 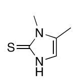 | 15 ± 3.5 | NT |
| **57** | 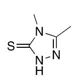 | >100 | NT |
| **58** | 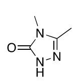 | >100 | NT |
| **59** | 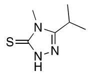 | 7.7 ± 1.0 | NT |

**Supplementary Table 1.** Chemical structures of the initial hit from uHTS (**39**) and analogues purchased based on this hit compound. TR-FRET data are reported as the $IC_{50}$ ± standard deviation (n = 3). DSF data are reported as **Δ**Tm ± standard deviation (n = 3). [a]TR-FRET data from initial uHTS. NT = Not tested.

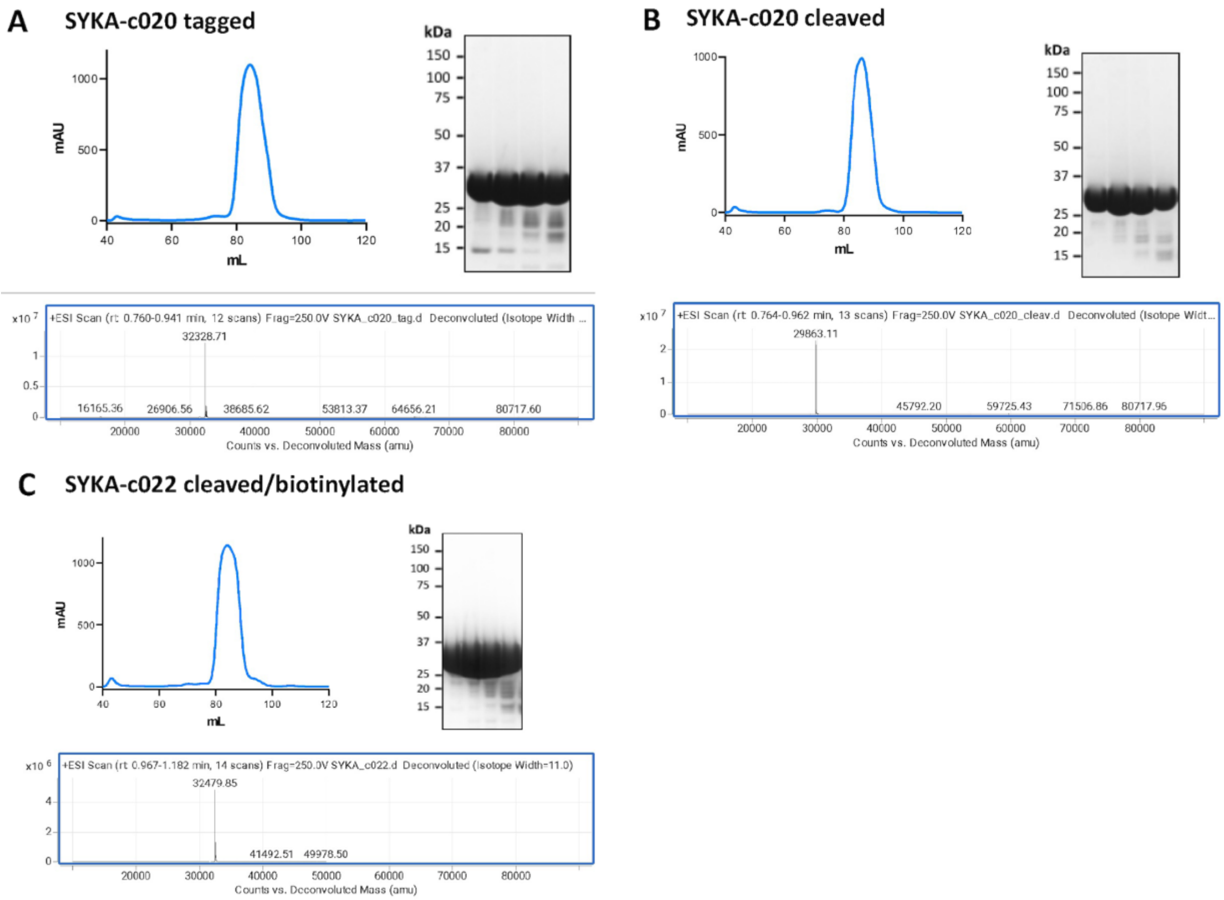

**Supplementary Figure 1.** Gel filtration fractions and intact mass deconvolution of purified SYK proteins: **(A)** SYKA-c020 His tagged, **(B)** SYKA-c020 His tag cleaved, and **(C)** SYKA-c020 His tag cleaved and biotinylated.

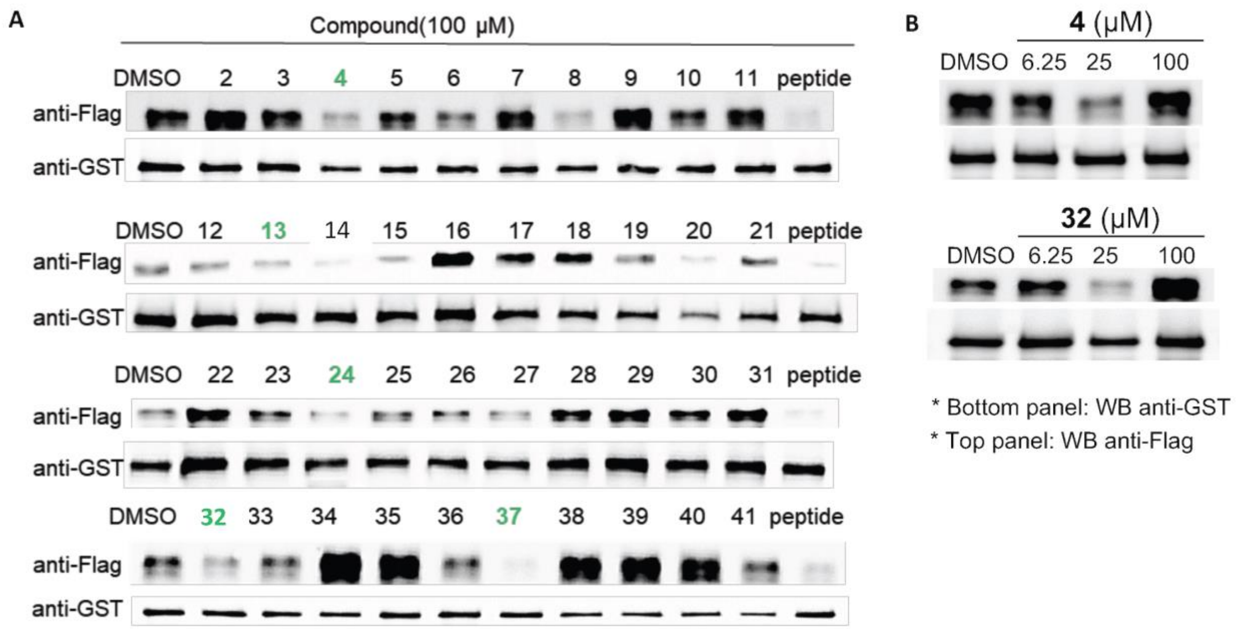

**Supplementary Figure 2.** Second biological replicate of GST-PD data displayed in Figure 4B and dose-response GST-PD of compounds **4** and **32. (A)** Compounds **4**, **13**, **24**, **32**, and **37**, **4**, were able to inhibit the interaction between SYK-GST and FCER1G-Flag in a pulldown assay at a single concentration (100 μM). Second biological replicate displayed. **(B)** Dose-response GST-PD of **4** and **32**.

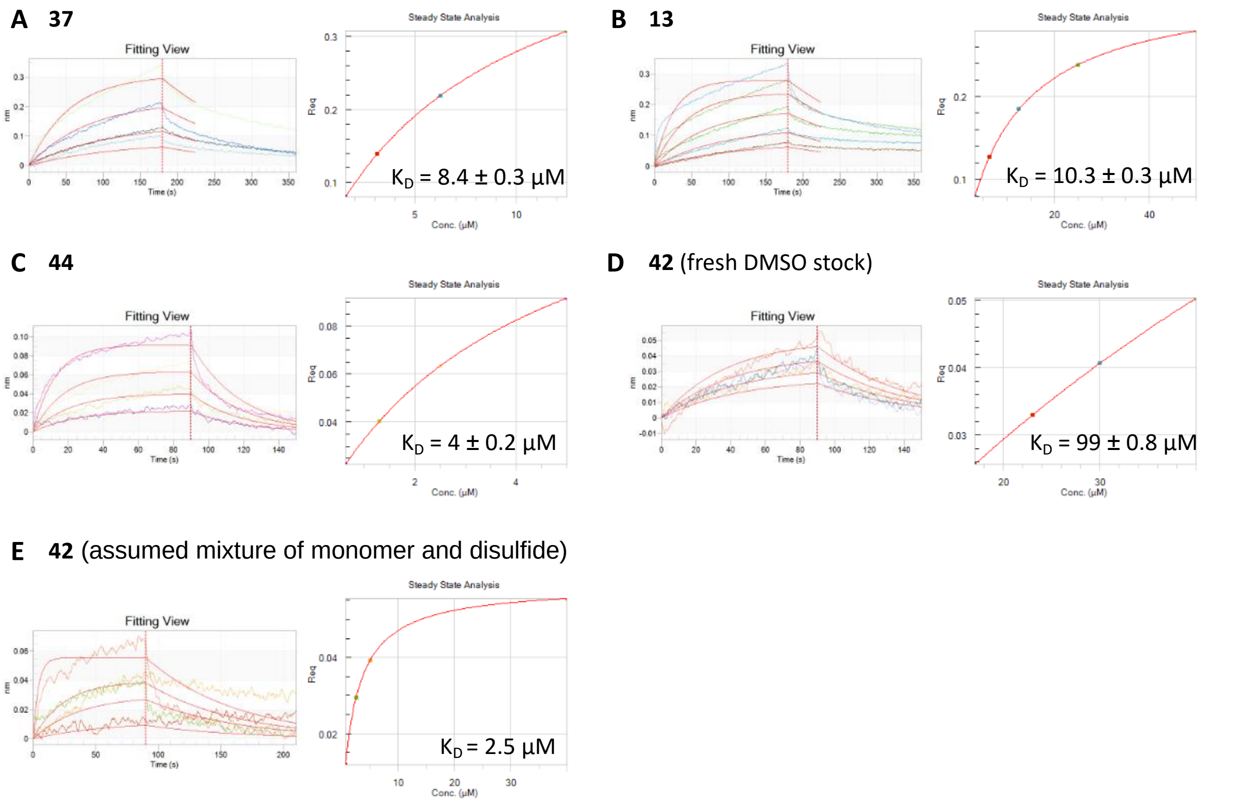

**Supplementary Figure 3.** Secondary biophysical analyses of hit compounds via BLI. **(A) 37**, **(B) 13**, **(C) 44**, **(D) 42** (fresh DMSO stock – monomer compound), **(E) 42** (assumed mixture of monomer and disulfide) BLI data.

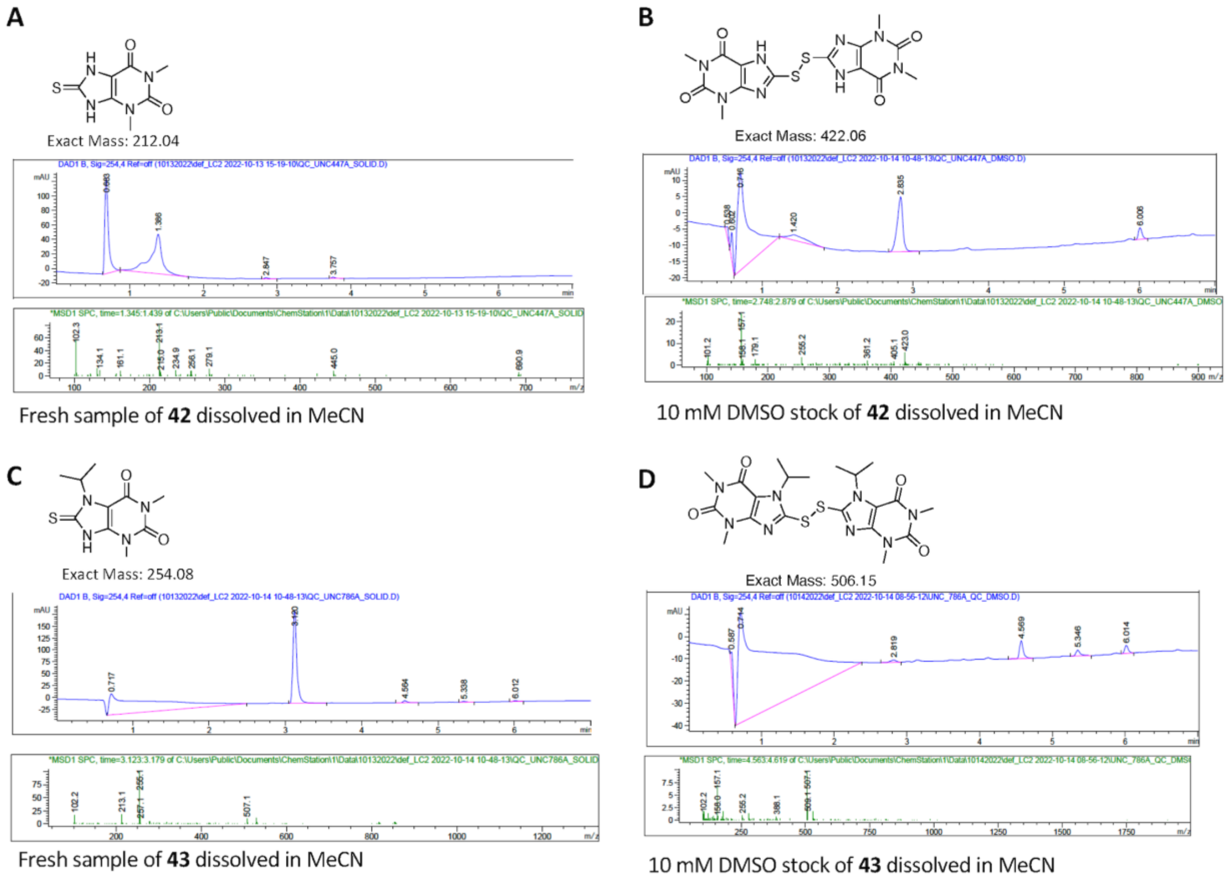

**Supplementary Figure 4.** Identification of an active disulfide compound which forms in DMSO solutions of **42** and **43**. **(A)** LCMS analysis of **42** as a fresh solution in MeCN indicates the monomer is present. LCMS Calculated for $[M+H]^+$ $C_7H_9N_4O_2S$: 213.04; observed: 213.1 $[M+H]^+$. **(B)** LCMS analysis of **42** 10 mM stock solution in MeCN indicates the disulfide dimer is present. LCMS Calculated for $[M+H]^+$ $C_{14}H_{15}N_8O_4S_2$: 423.06; observed: 423.0 $[M+H]^+$. **(C)** LCMS analysis of **43** as a fresh solution in MeCN indicates the monomer is present. LCMS Calculated for $[M+H]^+$ $C_{10}H_{15}N_4O_2S$: 255.08; observed: 255.1 $[M+H]^+$. **(D)** LCMS analysis of **43** 10 mM stock solution in MeCN indicates the disulfide dimer is present. LCMS Calculated for $[M+H]^+$ $C_{20}H_{27}N_8O_4S_2$: 507.15; observed: 507.1 $[M+H]^+$.

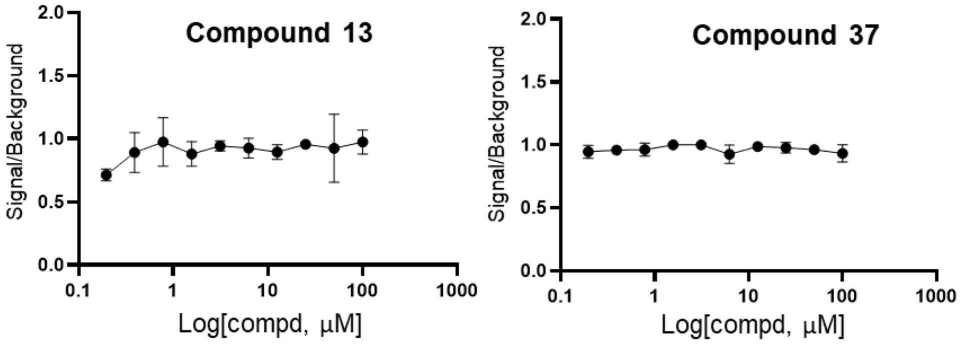

**Supplementary Figure S5.** Measurements of solubility of Compound **13** and **37** in PBS by Nephelometer.

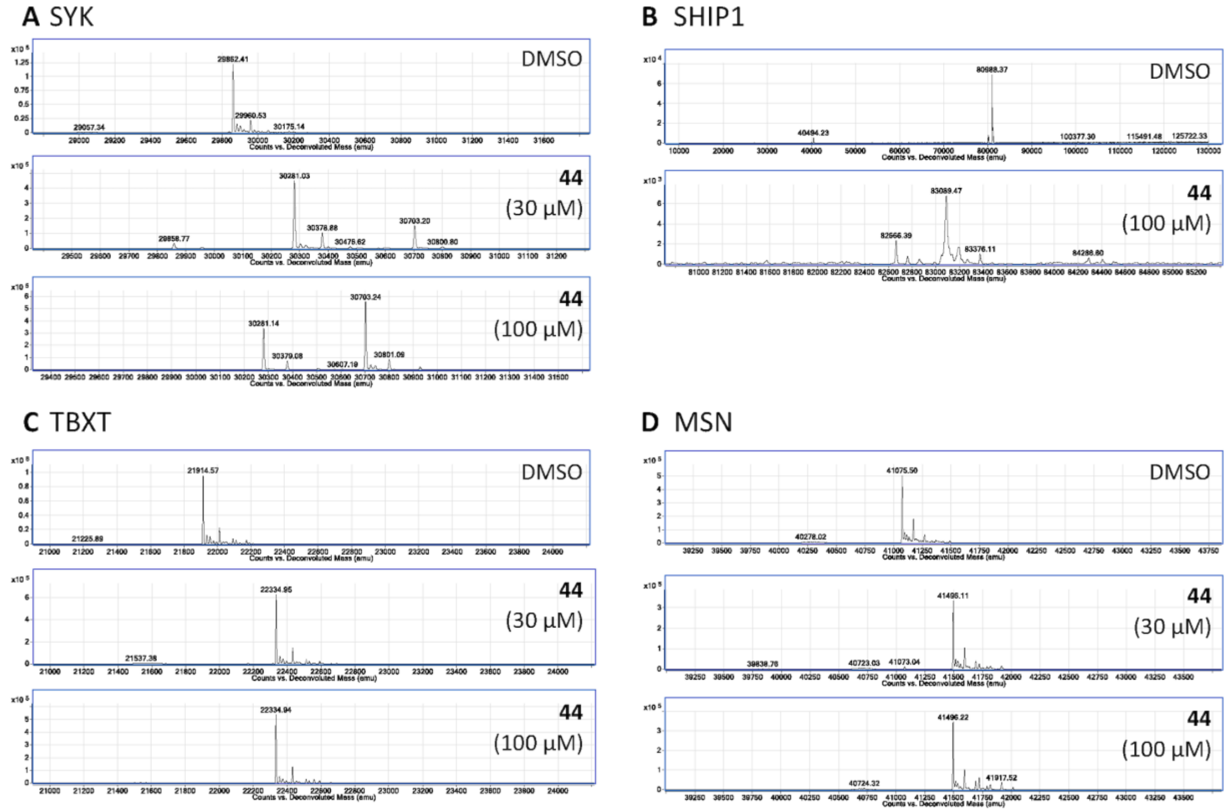

**Supplementary Figure 6.** Mass spectrometry of **44** incubated at 30 and 100 µM at rt for 1 h with proteins; **(A)** SYK, **(B)** SHIP1, **(C)** TBXT, **(D)** MSN.

### Chemistry

#### $^1$H NMR of 8,8'-disulfanediylbis(1,3-dimethyl-3,7-dihydro-1*H*-purine-2,6-dione) (44)

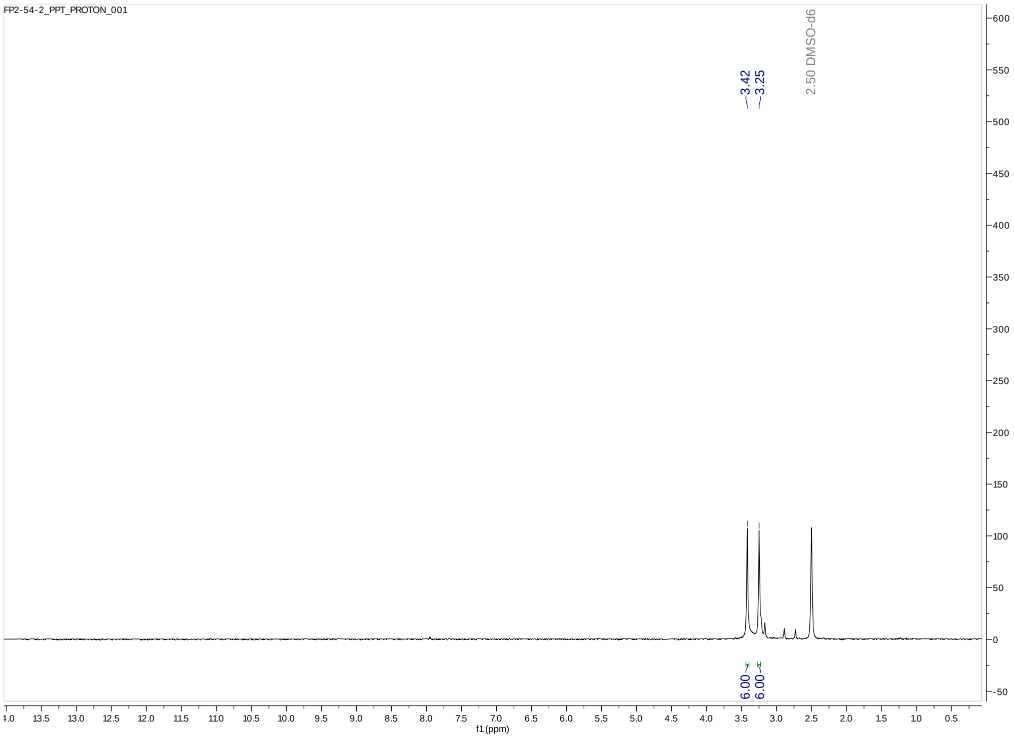

#### LCMS of 8,8'-disulfanediylbis(1,3-dimethyl-3,7-dihydro-1*H*-purine-2,6-dione) (44)

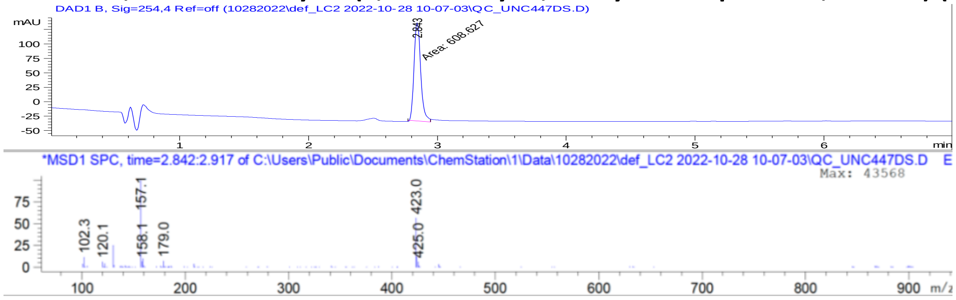

### Members of Emory-Sage-SGC TREAT-AD Center

Ishita Ajith[1], Joel K. Annor-Gyamfi[2], Jeff Aube[2], Alison D. Axtman[2], Frances M. Bashore[2], Ranjita S. Betarbet[3], Juan Botas[4], William J. Bradshaw[1], Paul E. Brennan[1], Peter J. Brown[2], Robert R. Butler 3rd[5], Jacob L. Capener[2], Gregory W. Carter[6], Gregory A. Cary[6], Catherine Chen[4], Rachel Commander[3], Sabrina Daglish[2], Suzanne Doolen[7], Yuhong Du[3], Aled M. Edwards[8], Michelle E. Etoundi[4], Kevin J. Frankowski[2], Stephen V. Frye[2], Haian Fu[3], Opher Gileadi[1], Marta Glavatshikh[2], Jake Gockley[9], Katerina Gospodinova[1], Anna K. Greenwood[9], Peter A. Greer[10], Lea T. Grinberg[11], Shiva Guduru[2], Levon Halabelian[8], Crystal Han[5], Brian Hardy[2], Laura M. Heath[9], Stephanie Howell[2], Andrey A. Ivanov[3], Suman Jayadev[12], Vittorio L. Katis[1], Stephen Keegan[6], May Khanna[13], Dmitri Kireev[2], Carl LaFlamme[14], Karina Leal[9], Tom V. Lee[4], Tina M. Leisner[2], Allan I. Levey[3], Qianjin Li[3], David Li-Kroeger[4], Zhandong Liu[4], Benjamin A. Logsdon[9], Frank M. Longo[5], Lara M. Mangravite[9], Peter S. McPherson[14], Richard M. Nwakamma[3], Felix O. Nwogbo[2], Carolyn

A. Paisie[6], Arti Parihar[12], Kenneth H. Pearce[2], Kun Qian[3], Min Qui[3], Stacey J Sukoff Rizzo[7], Karolina A. Rygiel[1], Julie Schumacher[5], David D. Scott[15], Nicholas T. Seyfried[3], Joshua M. Shulman[4], Ben Siciliano[3], Arunima Sikdar[2], Nathaniel Smith[4], Michael Stashko[2], Judith A. Tello Vega[15], Dilipkumar Uredi[2], Dongxue Wang[3], Jianjun Wang[3], Xiaodong Wang[2], Zhexing Wen[3], Jesse C. Wiley[9], Alexander Wilkes[1], Charles A. Williams[12], Timothy M. Willson[2], Aliza Wingo[3], Thomas S. Wingo[3], Novak Yang[3], Jessica E. Young[12], Miao Yu[6], Elizabeth L. Zoeller[3]
